# Multiple domains in ARHGAP36 regulate PKA degradation and Gli activation

**DOI:** 10.1101/2020.05.14.094961

**Authors:** Patricia R. Nano, Takamasa Kudo, Nancie A. Mooney, Jun Ni, Janos Demeter, Peter K. Jackson, James K. Chen

## Abstract

ARHGAP36 is a Rho GTPase-activating protein (GAP) family member that contributes to spinal cord development and tumorigenesis. This multidomain protein is composed of splicing-dependent N-terminal sequences, the GAP-like region, and a unique C-terminal domain, and an N-terminal arginine-rich region has been shown to suppress protein kinase A (PKA) and activate Gli transcription factors. To understand how these structural elements act in concert, we have mapped the ARHGAP36 structure-activity landscape with domain- and amino-acid-level resolution. ARHGAP36-mediated Gli activation can be repressed by N-terminal sequences that regulate subcellular ARHGAP36 localization and PKA targeting. The GAP-like and C-terminal domains counteract this autoinhibitory mechanism and promote ARHGAP36 trafficking to the plasma membrane and primary cilium, respectively. The GAP-like domain may also conditionally suppress the arginine-rich region, and it modulates ARHGAP36 binding to the prolyl oligopeptidase-like protein PREPL and the E3 ubiquitin ligase PRAJA2. These domain-dependent activities provide a potential means for tissue-specific ARHGAP36 functions.

## INTRODUCTION

Gli transcription factors (Gli1-3) are essential regulators of cell proliferation and differentiation, controlling fate specification in the neural tube (*Briscoe, et al., 2000; Stamataki, et al., 2005*) and limb bud (*Hill, et al., 2009; te Welscher, et al., 2002*) and the maintenance of granule neuron precursors in the developing cerebellum (*Lewis, et al., 2004; Wallace, 1999; Wechsler-Reya, et al., 2001*). Accordingly, misregulation of Gli activity can lead to uncontrolled cell growth, resulting in basal cell carcinoma, medulloblastoma, and other human cancers (*Hui, et al., 2011*). Gli functions are primarily regulated by the Hedgehog (Hh) pathway, and in the absence of Hh ligands, GLI2 and GLI3 are bound to the scaffolding protein Suppressor of Fused (SUFU) (*Stone, et al., 1999; Wang, Chengbing, et al., 2010*), which promotes their sequential phosphorylation by protein kinase A (PKA), glycogen synthase kinase β (GSK3β), and casein kinase 1 (CK1) (*Pan, et al., 2006; Pan, et al., 2007; Tempe, et al., 2006; Wang, Baolin, et al., 2006*). Proteasomal machinery is then recruited to the phosphorylated Gli proteins, resulting in GLI2 degradation and proteolytic conversion of GLI3 into a transcriptional repressor (*Pan, et al., 2007*).

Hh ligands suppress these intracellular processes, acting via the transmembrane receptors Patched1 (PTCH1) and Smoothened (SMO). Through mechanisms that remain unclear, SMO promotes the dissociation of GLI2 and GLI3 from SUFU, uncoupling the transcription factors from proteasomal regulation and allowing the full-length proteins to become transcriptional activators (*Humke, et al., 2010; Tukachinsky, et al., 2010*). SMO activity is suppressed by PTCH1 (*Murone, et al., 1999; Rohatgi, et al., 2007; Taipale, J., et al., 2002*), which is in turn directly inhibited by Hh ligands (*Incardona, et al., 2000; Stone, et al., 1996*). Hh signaling therefore induces SMO activation and the expression of Hh target genes, including those that encode PTCH1 (*Ågren, et al., 2004*) and the constitutively active transcription factor GLI1 (*Bai, et al., 2004; Dai, et al., 1999*). The primary cilium serves a key center for these signaling events (*Dorn, et al., 2012; Haycraft, et al., 2005; Kim, et al., 2009; May, et al., 2005; Rohatgi, et al., 2007; Wang, Yu, et al., 2009; Wen, et al., 2010*), and this cell-surface protrusion is required for both Gli activator and repressor formation (*Huangfu, et al., 2005; Liu, Aimin, et al., 2005*).

In addition to these signal transduction mechanisms, there is growing evidence for non-canonical Gli regulation in both normal physiology (*Dennler, et al., 2007; Flora, et al., 2009; Riobó, et al., 2006*) and in cancer (*Beauchamp, et al., 2009; Dennler, et al., 2007; Elsawa, et al., 2011; Han, et al., 2015; Kasper, et al., 2006; Liu, Z., et al., 2014; Long, et al., 2014*). Our laboratory previously established ARHGAP36 as a non-canonical Gli activator that acts in a SMO- independent manner (*Rack, et al., 2014*). We identified this Rho GTPase-activating protein (GAP) family member in a genome-scale screen for Hh pathway agonists (*Rack, et al., 2014*), and subsequent studies have uncovered an essential role for ARHGAP36 in the specification of lateral motor column neurons (*Nam, et al., 2019; Rack, et al., 2014*). Endogenous *Arhgap36* transcription in the developing mouse spinal cord coincides with Hh pathway activation, and its overexpression leads to ectopic induction of the Hh target genes *Ptch1* and *Gli1* (*Nam, et al., 2019*). In addition, *Arhgap36* expression has been found to correlate with SMO inhibitor resistance in Hh pathway-driven murine medulloblastomas (*Buonamici, et al., 2010; Rack, et al., 2014*), and upregulating this Rho GAP family member in neural progenitor cells is sufficient to induce medulloblastomas in mice (*Beckmann, et al., 2019*). ARHGAP36 may promote tumor growth through multiple mechanisms, as elevated *ARHGAP36* expression also has been associated with Hh pathway-independent subtypes of medulloblastoma and neuroblastoma (*Beckmann, et al., 2019; Lee, et al., 2019*).

Despite the emerging importance of ARHGAP36 in neuronal development and cancer, the biochemical and cellular mechanisms that regulate and transduce its activity are not well understood. The ARHGAP36 protein consists of unique N- and C-terminal domains and a central region that is homologous to Rho GAPs. In addition, the five annotated isoforms of human ARHGAP36 have varying N-terminal structures due to alternative splicing (Figure 1A). Four of the ARHGAP36 isoforms can activate Gli proteins, with the gene product harboring the longest N-terminal domain (isoform 1) being the sole exception (*Rack, et al., 2014*). Isoform 1 is also the only ARHGAP36 protein that does not localize to the plasma membrane, and it instead adopts a perinuclear distribution (*Müller, et al., 2020; Rack, et al., 2014*). In addition, the shortest ARHGAP36 protein (isoform 3) accumulates in the primary cilium, whereas other ARHGAP36 isoforms cannot be detected in this signaling center under steady-state conditions (*Rack, et al., 2014*). More recently, it has been shown that an N-terminal arginine-rich motif conserved in all human ARHGAP36 isoforms can bind directly to catalytic subunits of PKA (PRKACA and PRKACB; henceforth referred to as PKA_cat_) (*Eccles, et al., 2016*). In the context of isoform 2, this motif mediates the degradation of PKA_cat_, and a 77-amino-acid N-terminal fragment that includes this arginine-rich region has been shown to be necessary and sufficient for cellular PKA_cat_ depletion (*Eccles, et al., 2016*). These findings suggest a role for N-terminal sequences in targeting ARHGAP36 to specific subcellular compartments and establish PKA_cat_ inhibition as a potential basis for ARHGAP36-mediated Gli activation.

**Figure 1.**
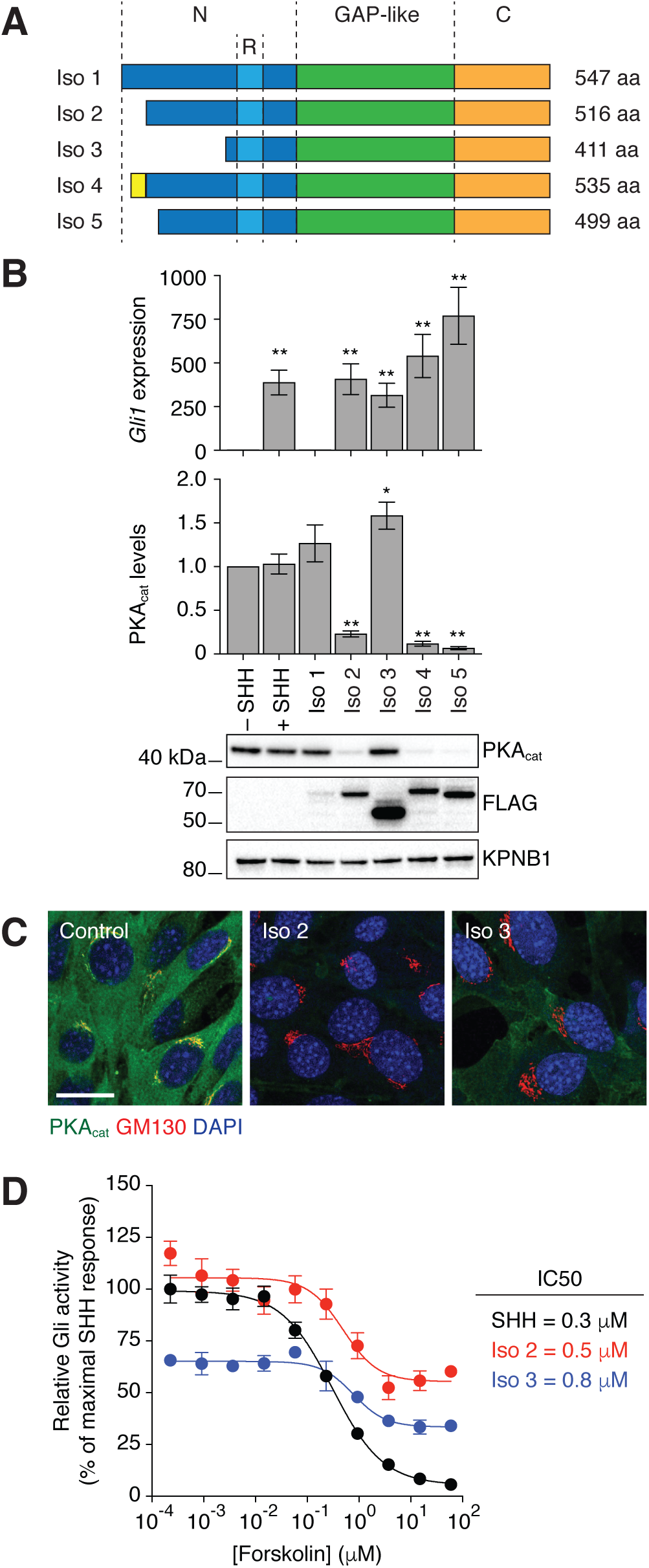
ARHGAP36 isoforms differentially induce PKAcat depletion and Gli activation. (A) Domain architecture of the five human ARHGAP36 isoforms, with the N-terminal domain (N) shown in dark blue, arginine-rich motif (R) in light blue, GAP-like domain in green, and C-terminal domain (C) in orange. The yellow region indicates an amino acid sequence unique to isoform 4. (B) *Gli1* mRNA and PKAcat protein levels in NIH-3T3 cells retrovirally transduced with the indicated FLAG-tagged ARHGAP36 isoforms. Uninfected cells treated with or without Sonic Hedgehog (SHH) ligand were included as positive and negative controls, respectively. Data are the average fold change relative to the negative control for three biological replicates ± s.e.m. Single and double asterisks indicate *P* < 0.05 and *P* < 0.01, respectively. A representative western blot for each condition is also shown. (C) PKAcat localization in NIH-3T3 cells transduced with FLAG-tagged ARHGAP36 isoform 2 or 3. Representative immunofluorescence micrographs are shown with staining for PKAcat, GM130 (cis-Golgi), and DAPI (nucleus). Scale bar: 20 μm. (D) Forskolin dose-response curves for SHH-LIGHT2 cells stimulated with SHH or transduced with FLAG-tagged ARHGAP36 isoform 2 or 3. Data are the average Gli reporter activities for at least three biological replicates ± s.e.m., normalized to the maximum response in SHH-treated cells.

In comparison, the functions of the GAP-like and C-terminal domains in ARHGAP36 have not yet been elucidated. Rho GAP family members typically attenuate the function of Rho GTPases by stimulating GTP hydrolysis (*Moon, 2003*). However, the GAP-like region in ARHGAP36 lacks the “arginine finger” motif conserved in catalytically active homologs (*Rack, et al., 2014; Scheffzek, et al., 1998*), and ARHGAP36 has no effect on the activities of Rac1, Cdc42, and Rho A (*Müller, et al., 2020*). In addition, ARHGAP36 residues that are structurally equivalent to those previously associated with Rho GAP-catalyzed GTP hydrolysis are not required for ARHGAP36-mediated Gli activation (*Rack, et al., 2014*). Non-catalytic mechanisms have been reported for several Rho GAP family members (*Amin, et al., 2016; Faucherre, et al., 2003; Marchesi, et al., 2014*), and it is possible that the GAP-like domain in ARHGAP36 similarly interacts with Rho GTPases or other signaling proteins in a stoichiometric manner. How the C-terminal domain might contribute to ARHGAP36 function is even more enigmatic since it lacks sequence homology with other proteins.

Deciphering the molecular and cellular mechanisms that regulate ARHGAP36 activity requires a deeper understanding of the relationship between ARHGAP36 structure and function. Here we describe our systematic mapping of the ARHGAP36 structure-activity landscape using individual ARHGAP36 isoforms, truncated variants, and a high-throughput mutagenesis screen. Our findings demonstrate that ARHGAP36-dependent Gli activation and cellular PKA_cat_ depletion are separable activities and reveal isoform-specific differences in subcellular PKA_cat_ targeting. While the ARHGAP36 N-terminal domain is necessary and sufficient for Gli activation, an N-terminal region in isoform 2 (residues 1-105; N2_1-105_) can inhibit this function and suppress protein localization to the plasma membrane. This autoinhibitory mechanism is counteracted by the GAP-like and C-terminal domains, which promote ARHGAP36 recruitment to the plasma membrane and primary cilium, respectively. Finally, we have discovered several residues within the GAP-like domain that are necessary for Gli activation by full-length ARHGAP36 isoforms. These residues are predicted to cluster within the GAP-like domain structure, at a site distal to the Rho GTPase-binding pocket, and they are required for ARHGAP36 recruitment to the plasma membrane. We have also leveraged these mutants to discover factors that bind specifically to the wild-type protein, identifying potential mediators of ARHGAP36 function. Taken together, our work supports a model in which ARHGAP36 activity state, subcellular localization, and effector binding are regulated by structural elements distributed throughout protein. In combination with the differential expression of ARHGAP36 isoforms, such mechanisms could allow ARHGAP36 to control Gli activity and other PKA_cat_-regulated processes in a tissue-specific manner.

## RESULTS

### Gli activation and cellular PKA_cat_ depletion are separable ARHGAP36 functions

To explore the relationship between ARHGAP36-mediated Gli activation and PKA_cat_ degradation, we measured the effect of each human ARHGAP36 isoform on both cellular processes. Individual isoforms were retrovirally transduced into NIH-3T3 mouse fibroblasts, a commonly used line for studying Hh signal transduction (*Taipale, J, et al., 2000*) that also exhibits ARHGAP36 responsiveness (*Eccles, et al., 2016; Rack, et al., 2014*). The resulting levels of *Gli1* mRNA and PKA_cat_ protein were assessed by qRT-PCR and western blot, respectively. Overexpression of isoforms 2, 4, or 5 was sufficient to activate Gli and deplete the cells of PKA_cat_, while isoform 1 exhibited neither activity (Figure 1B). In contrast, transduction of isoform 3 induced *Gli1* expression without reducing cellular PKA_cat_ levels to a discernable extent. These results indicate that total PKA_cat_ depletion is not required for ARHGAP36-mediated Gli activation, raising the possibility that ARHGAP36 regulates Gli proteins by targeting a specific subcellular pool of PKA_cat_ and/or through PKA-independent mechanisms.

We investigated these two models by further comparing the activities of ARHGAP36 isoform 3 with those of isoform 2. Using immunofluorescence microscopy, we observed that isoform 2 globally depleted PKA_cat_ in NIH-3T3 cells, corroborating our western blot analyses (Figure 1C). In contrast, isoform 3 reduced PKA_cat_ pools predominantly in the Golgi. These findings are consistent with the accumulation of isoform 3 in the primary cilium (*Rack, et al., 2014*), as the cilium base communicates directly with the Golgi through vesicular trafficking (*Pedersen, et al., 2016*). We next examined how the activities of ARHGAP36 isoforms 2 and 3 are affected by forskolin, an adenylate cyclase agonist that increases cAMP levels and PKA_cat_ activity. We transduced the ARHGAP36 constructs into NIH-3T3 fibroblasts stably expressing a Gli-dependent firefly luciferase reporter (SHH-LIGHT2 cells) (*Taipale, J, et al., 2000*) and treated the cells with the PKA_cat_ activator. Although the isoform 2-expressing cells exhibited almost two-fold higher Gli reporter activity than those expressing isoform 3, forskolin inhibited Gli function in both lines with comparable IC50s (Figure 1D). Maximal doses of the PKA_cat_ activator also suppressed about 50% of the Gli reporter activity induced by each isoform. Together, these results demonstrate that N-terminal sequences in ARHGAP36 can regulate its ability to target PKA_cat_ in specific subcellular compartments and raise the possibility that ARHGAP36 activates Gli proteins through both PKA_cat_-dependent and -independent mechanisms.

### ARHGAP36 is autoinhibited by an N-terminal region that is counteracted by the GAP-like and C-terminal domains

We continued to investigate functional differences between ARHGAP36 isoforms 2 and 3 by determining the activities of various truncation mutants (Figure 2A). By retrovirally expressing these constructs in NIH-3T3 cells, we observed that the N-terminal domain of ARHGAP36 isoform 2 (residues 1-194; N2) is necessary and sufficient for its effects on Gli and PKA_cat_ (Figure 2B) corroborating previous reports (*Eccles, et al., 2016*). However, N2 was less effective at activating *Gli1* expression than the N-terminal domain of isoform 3 (N3), even though N2 could induce PKA_cat_ degradation. N2 was also markedly less active than full-length isoform 2. In contrast, N3 and full-length isoform 3 could induce *Gli1* expression to similar extents (Figure 2B). These results indicate that the N2 region absent in isoform 3 (residues 1-105; N2_1-105_) represses the Gli-activating function of the remaining N-terminal domain. Moreover, our findings suggest that ARHGAP36 sequences in the GAP-like and/or C-terminal domains can influence N2_1-105_ function.

**Figure 2.**
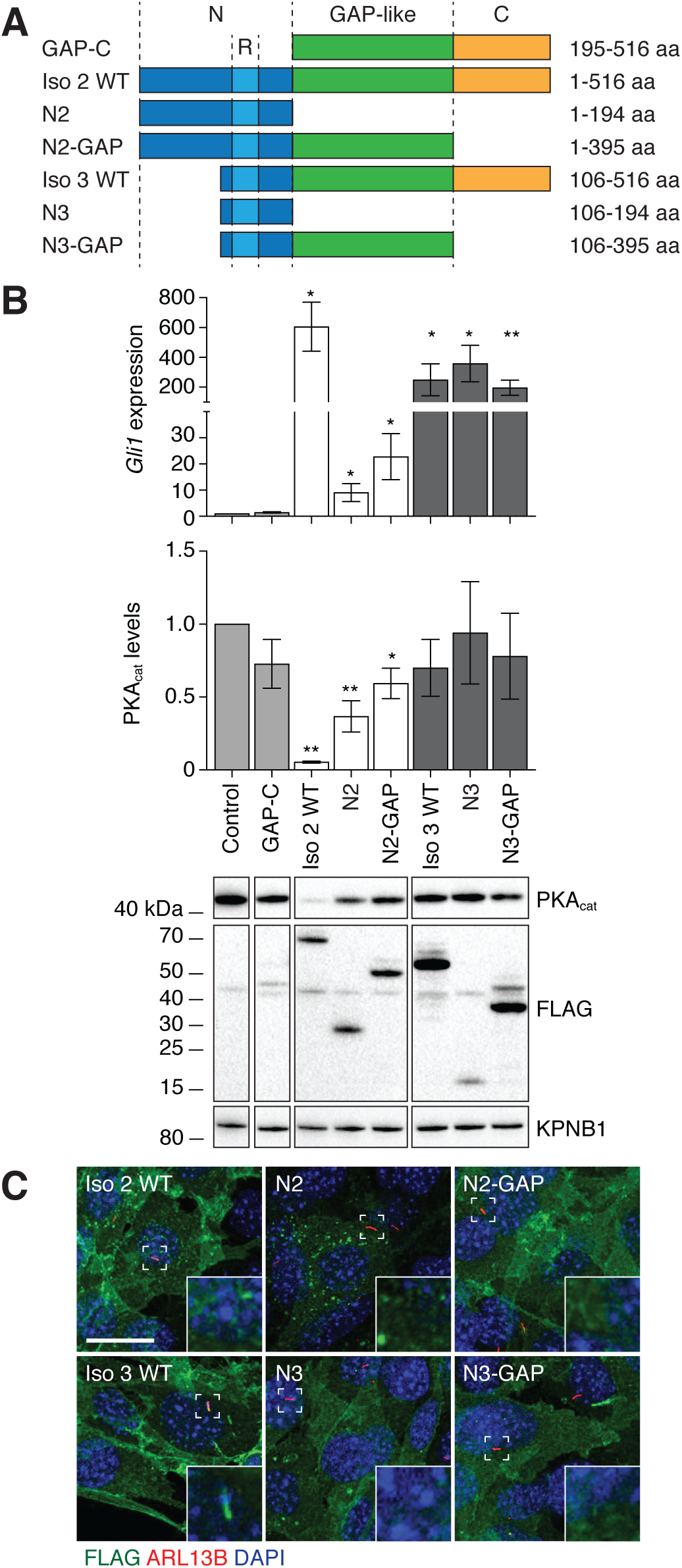
N-terminal, GAP-like, and C-terminal domains have opposing effects on ARHGAP36 function. (A) Schematic representation of ARHGAP36 isoform 2 or 3 truncation mutants. Residue numbers are based on the amino acid sequence of isoform 2. (B) *Gli1* mRNA and PKAcat protein levels in NIH-3T3 cell retrovirally transduced with the indicated FLAG-tagged ARHGAP36 truncation mutants. Data are the average fold change relative to uninfected cells for three biological replicates ± s.e.m. Single and double asterisks indicate *P* < 0.05 and *P* < 0.01, respectively. A representative western blot for each condition is also shown (lanes from the same blot image have been cropped and re-ordered for clarity). (C) Subcellular distributions of the indicated FLAG-tagged ARHGAP36 constructs in NIH-3T3 cells. Representative immunofluorescence micrographs are shown with staining for FLAG, ARL13B (primary cilium) and DAPI (nucleus). Insets highlight ciliated regions in the dashed boxes. Scale bar: 20 μm. Images were processed to establish comparable maximum pixel intensities in order to highlight differences in localization.

To discern how the GAP-like and C-terminal domains contribute to ARHGAP36 function, we examined the activities of N2-GAP and N3-GAP constructs in NIH-3T3 cells. N2-GAP was moderately more active than N2 with respect to *Gli1* expression, but it was still much less active than full-length isoform 2 (Figure 2B). N3-GAP activity was comparable to that of N3 and full-length isoform 3. Thus, both the GAP-like and C-terminal domains can counteract the autoinhibitory function of N2_1-105_, with the C-terminal region playing a particularly important role. In the absence of N2_1-105_, ARHGAP36 does not require its GAP-like and C-terminal domains to achieve high levels of Gli activity.

We next sought to determine how the N2_1-105_ region, GAP-like domain, and C-terminal domain regulate ARHGAP36 function. The differing subcellular distributions of full-length isoforms 2 and 3 indicate that the N2_1-105_ region influences ARHGAP36 trafficking (*Rack, et al., 2014*), and we therefore examined the localizations of the truncation mutants. N2 accumulated in both punctate structures and the plasma membrane, whereas N2-GAP was robustly recruited to the latter (Figure 2C). In comparison, both N3 and N3-GAP predominantly associated with the plasma membrane, and unlike the full-length isoform 3, neither construct accumulated in the primary cilium.

These domain-dependent changes in protein localization indicate that N2_1-105_ impedes and the GAP-like domain facilitates ARHGAP36 translocation to the plasma membrane. In principle, these opposing activities could involve direct interactions between the two regions in ARHGAP36 or parallel functions involving other factors. In addition, the C-terminal domain acts independently of the N2_1-105_ region to promote ciliary accumulation of ARHGAP36. The disparate roles of these domains in ARHGAP36 trafficking correlate with their divergent effects on Gli activation, providing further evidence that ARHGAP36 targets specific subcellular compartments to regulate Gli proteins.

### Identification of ARHGAP36 residues that are required for Gli activation

To characterize these regulatory ARHGAP36 sequences with amino-acid resolution, we developed a high-throughput mutagenesis screen for ARHGAP36 residues that are essential for Gli activation (Figure 3A). We used error-prone PCR to create a library of ARHGAP36 isoform 2 variants C-terminally fused to mCherry, obtaining a collection of approximately 100,000 single point mutants (27% of all library constructs and 22-fold theoretical coverage of the 4,374 possible variants). The library was retrovirally transduced into NIH-3T3 fibroblasts expressing a Gli-dependent green-fluorescent reporter (SHH-EGFP cells) (*Hyman, et al., 2009*) using a multiplicity of infection (MOI) of 0.3 to maximize the number of cells with single integration events. Cells expressing full-length ARHGAP36 proteins were then isolated by fluorescence-activated cell sorting (FACS) according to their mCherry fluorescence. We next cultured these cells under Hh signaling-competent conditions to allow active ARHGAP36 mutants to induce Gli-dependent EGFP expression, after which mCherry+/EGFP– cells were isolated by FACS. To ensure that these ARHGAP36-expressing cells still harbored a functional EGFP reporter, they were subsequently cultured with the SMO agonist SAG (*Chen, et al., 2002*). The resulting mCherry+/EGFP+ cells were obtained by FACS, yielding a population of cells expressing putative, inactive ARHGAP36-mCherry mutants.

**Figure 3.**
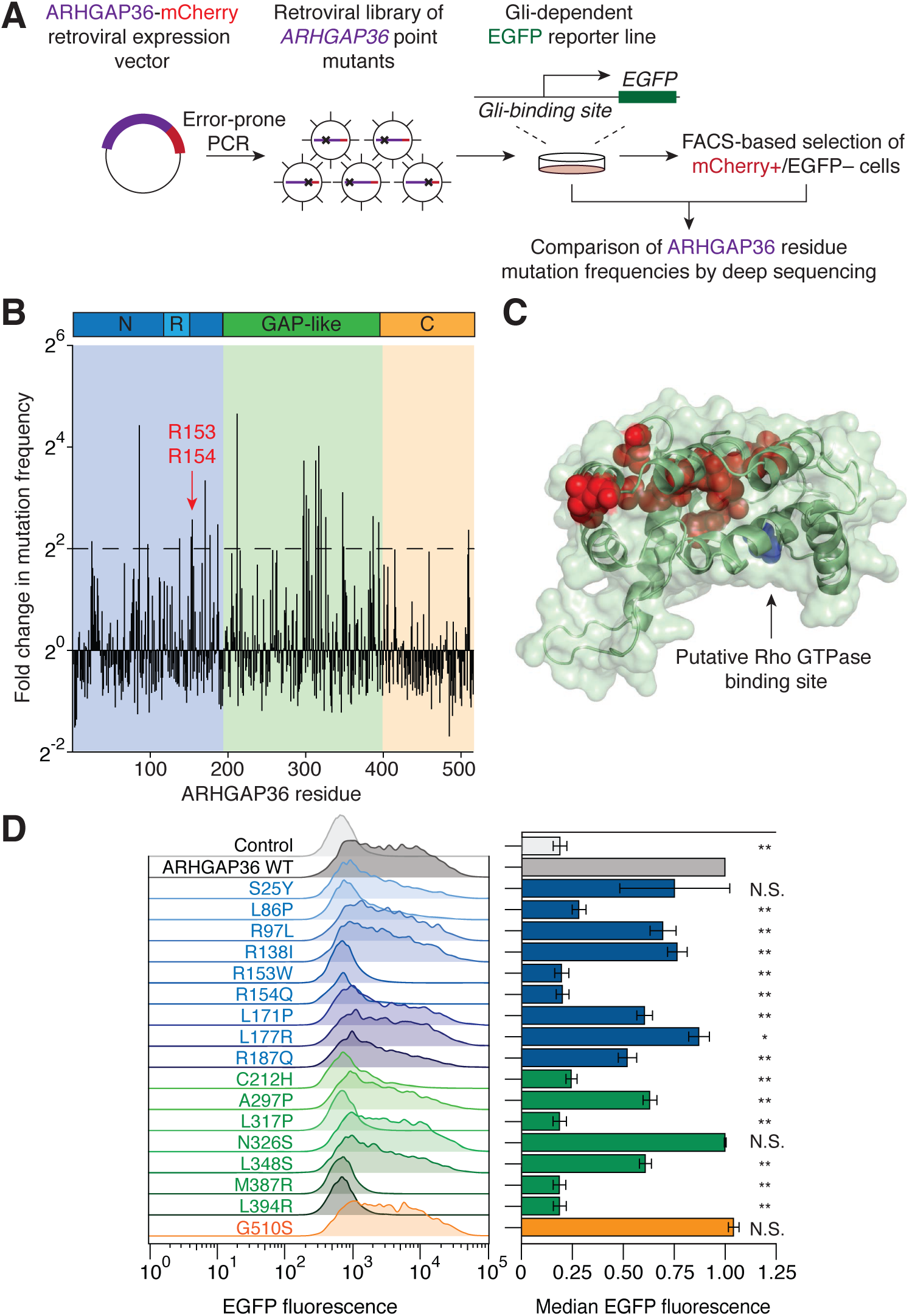
Identification of essential residues within the N-terminal and GAP-like domains. (A) Schematic representation of the high-throughput mutagenesis screen used to identify individual residues that contribute to ARHGAP36 function. (B) Histogram depict-ing the fold change in mutation frequency between pre- and post-selected population for each residue in ARHGAP36. (C) Homology model of the ARHGAP36 GAP-like domain structure based on the crystal structure of β2-chimaerin (PDB ID: 1xa6). Residues with > 4-fold change in mutation frequency are shown in red, and the site (Thr227) that is structurally equivalent to the arginine finger is shown in blue. (D) Activities of selected ARHGAP36 variants in SHH-EGFP cells, as assessed by flow cytometry-based measurements of Gli reporter fluorescence. The distributions (left) and relative medians (right) of EGFP fluorescence are shown for each ARHGAP36 construct. Data are the average fold change in median EGFP fluorescence relative to that of cells expressing wild-type ARHGAP36 for three biological replicates ± s.e.m. Single and double asterisks indicate *P* < 0.05 and *P* < 0.01, respectively.

To identify inactivating point mutations, we used genomic PCR and deep sequencing to compare the mutation frequency of each amino acid position in the pre- and post-selection populations. This analysis revealed several residues that could be required for Gli activation, including two N-terminal arginines (R153 and R154) that were previously shown to be required for ARHGAP36-mediated PKA_cat_ inhibition (Figure 3B, Supplementary File 1) (*Eccles, et al., 2016*). The majority of these putative essential residues were located in the GAP-like region, and structure homology modeling of this domain predicts that these amino acids cluster together at a site that is distal to the predicted Rho GTPase binding pocket (Figure 3C). We then used flow cytometry-based assays to validate a subset of these ARHGAP36 variants, prioritizing residues that were mutated > 4-fold more frequently in the inactive mutant pool. In these experiments, 17 individual ARHGAP36 mutants were retrovirally transduced into cells at an MOI of 0.3 to standardize their expression levels. With the exception of S25Y, N326S, and G510S, all of these point mutations significantly decreased ARHGAP36 activity in SHH-EGFP reporter cells (Figure 3D).

### GAP-like domain mutations disrupt ARHGAP36 recruitment to the plasma membrane

Guided by the results of our mutagenesis screen, we investigated how individual point mutations might regulate specific aspects of ARHGAP36 function. Our studies focused on the three residues with the greatest fold change in pre- and post-selection mutation frequencies: N-terminal domain residue L86 and two sites within the GAP-like domain, C212 and L317 (Figure 4A). We first examined how mutations at these sites affect the ability of ARHGAP36 isoform 2 to induce *Gli1* expression and PKA_cat_ degradation in NIH-3T3 cells. All three variants exhibited diminished *Gli1* expression and PKA_cat_ depletion, with the L317P mutation resulting in complete loss of both ARHGAP36-dependent activities (Figure 4B). We next determined the effects of these mutations on the subcellular localization of ARHGAP36 isoform 2. The L86P variant retained the ability to localize to the plasma membrane, but both mutations in the GAP-like domain rendered ARHGAP36 cytosolic (Figure 4C).

**Figure 4.**
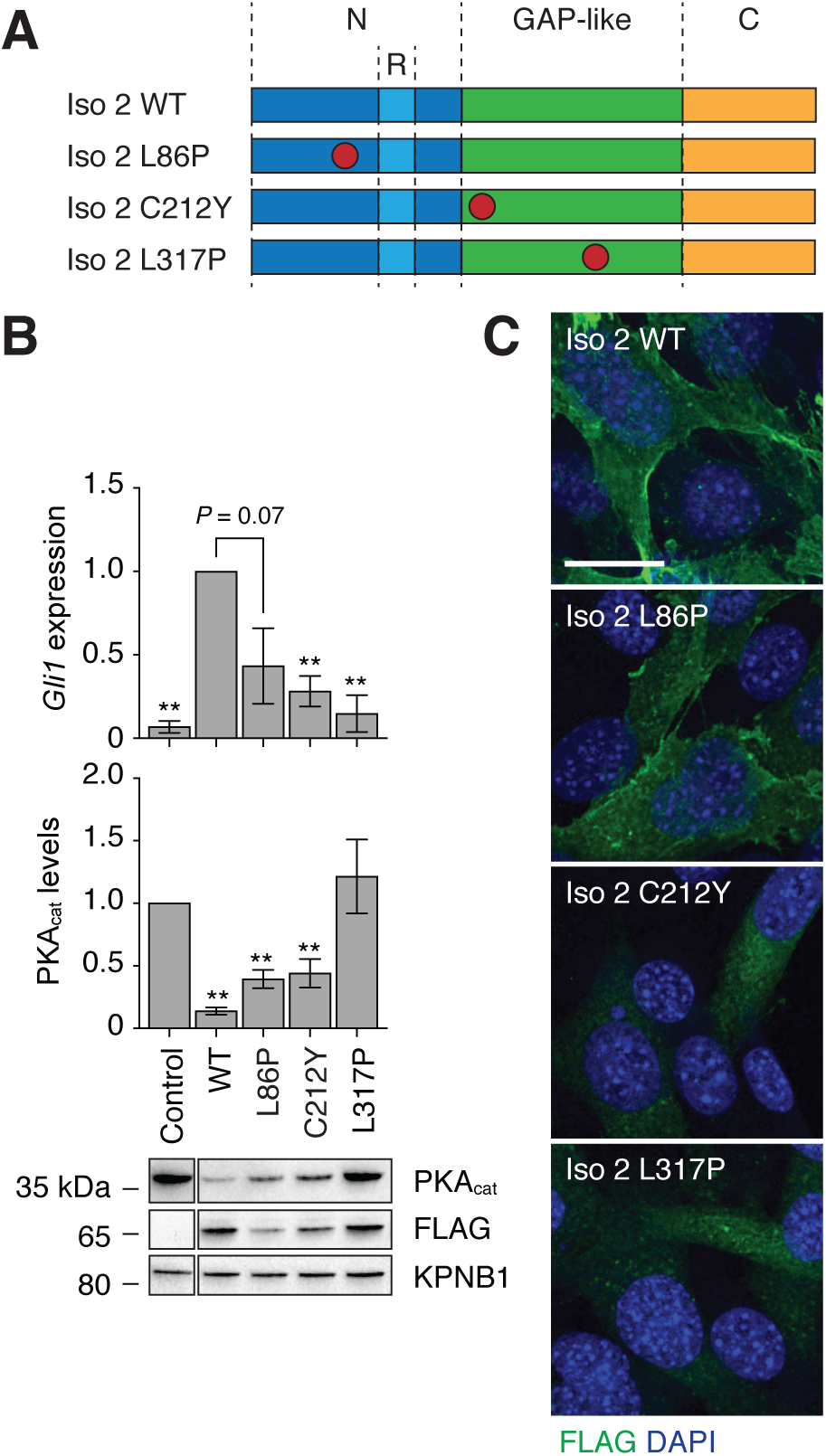
Point mutations in the GAP-like domain cause ARHGAP36 mislocalization. (A) Schematic representation of ARHGAP36 isoform 2 point mutants. (B) *Gli1* mRNA and PKAcat protein levels in NIH-3T3 cells retrovirally transduced with the indicated FLAG-tagged ARHGAP36 constructs. Uninfected cells were used as controls. Data are the average fold change relative to cells expressing wild-type ARHGAP36 (*Gli1* expression) or to untreated cells (PKAcat levels) for three biological replicates ± s.e.m. Single and double asterisks indicate *P* < 0.05 and *P* < 0.01, respectively. A representative western blot for each condition is also shown (lanes from the same blot image have been cropped and re-ordered for clarity). (C) Subcellular distributions of the indicated FLAG-tagged ARHGAP36 constructs in NIH-3T3 cells. Representative immunofluorescence micrographs are shown with staining for FLAG and DAPI (nucleus). Scale bar: 20 μm.

To further elaborate the contributions of individual residues to ARHGAP36 trafficking, we also assessed the localization of other ARHGAP36 point mutants that were inactive in SHH-EGFP cells (see Figure 3D). The R153W and R154Q variants accumulated in the plasma membrane, and the GAP-like domain point mutants M387R and L394R remained primarily in the cytosol (Figure 4 – figure supplement 1). Together, these results provide further evidence that the GAP- like domain recruits ARHGAP36 to the cell membrane, counteracting the function of N2_1-105_. We then investigated whether plasma membrane recruitment by the GAP-like domain requires N2_1-105_ or the C-terminal domain, both of which affect ARHGAP36 localization. As in the full-length isoform 2, C212Y and L317P mutations impeded the ability of N2-GAP to localize to the plasma membrane and activate Gli proteins (Figure 5A-B). Structurally equivalent mutations (C107Y and L212P) in ARHGAP36 isoform 3, which lacks the N2_1-105_ region, also rendered the protein cytosolic and attenuated Gli activation (Figure 5C-D). Thus, the two GAP-like domain mutations disrupt ARHGAP36 function independently of N2_1-105_ and the C-terminal domain.

**Figure 5.**
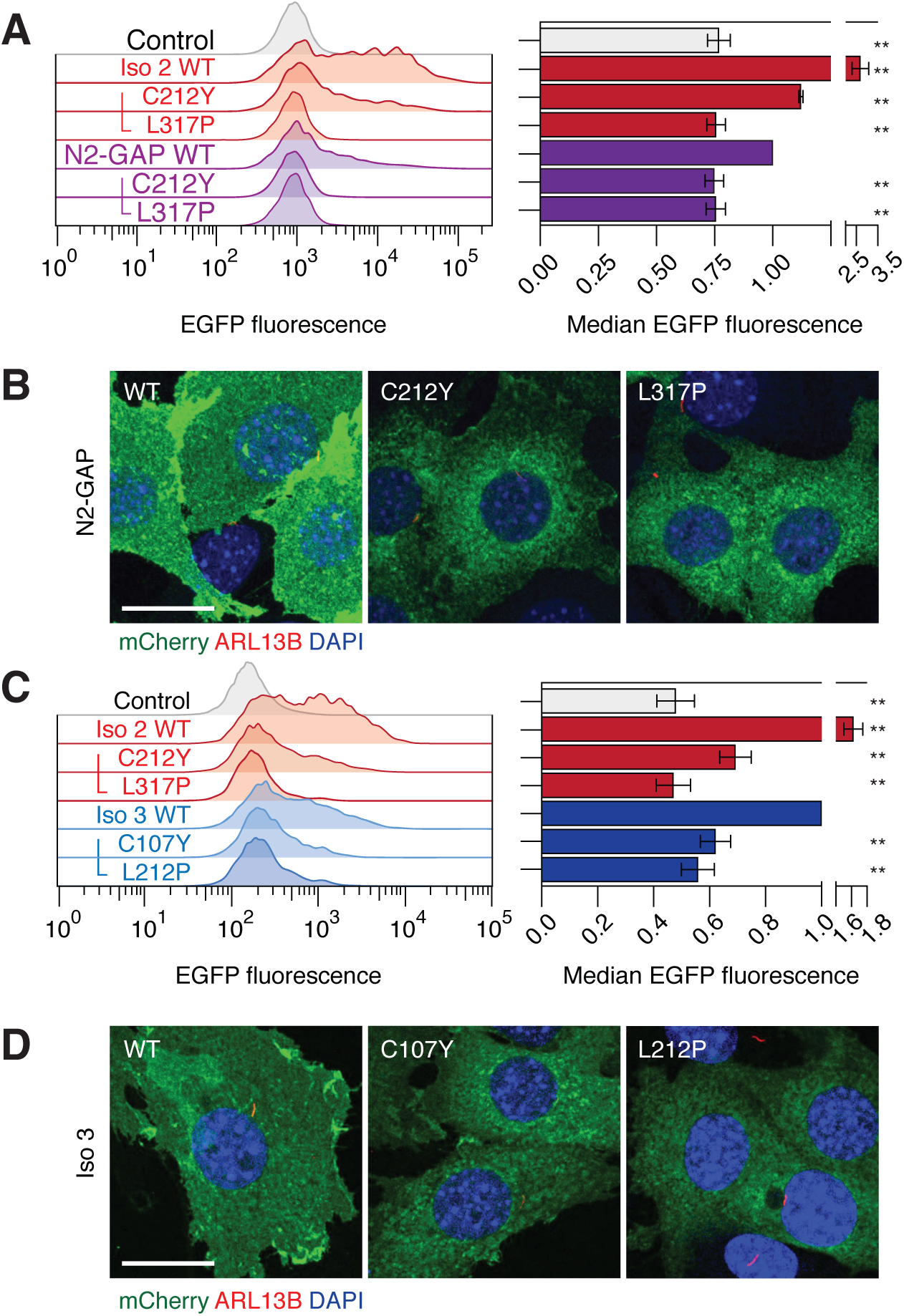
Point mutations in the GAP-like domain act independently of N21-105 and the C-terminal domain. (A and C) Effects of GAP-like domain point mutations in N2-GAP- or isoform 3-mediated Gli activation, as assessed in the flow cytometry-based SHH-EGFP assay. The distributions (left) and relative medians (right) of EGFP fluorescence are shown for each ARHGAP36 construct. Data are the average fold change in median EGFP fluorescence relative to that of cells expressing unmutated N2-GAP or isoform 3 for three biological replicates ± s.e.m. Double asterisks indicate *P* < 0.01. (B and D) Subcellular distributions of the indicated mCherry-tagged ARHGAP36 constructs in SHH-EGFP cells. Representative immunofluorescence micrographs are shown with staining for mCherry, ARL13B (primary cilium), and DAPI (nucleus). Scale bars: 20 μm.

### GAP-like domain mutations alter the ARHGAP36 interactome

In addition to uncovering functional roles for specific ARHGAP36 structures, our collection of inactive variants provided a means for identifying binding proteins that could participate in ARHGAP36-mediated Gli activation. Toward this goal, we compared the interactomes of wild-type and L317P ARHGAP36 isoform 2 by retrovirally transducing NIH-3T3 cells with vectors encoding each ARHGAP36 construct fused to a C-terminal LAP tag (S-peptide-PreScission protease site-EGFP) (*Ding, et al., 2016; Hsu, et al., 2019; Kanie, et al., 2017; Li, Bin, et al., 2017; Torres, et al., 2009; Wright, et al., 2011*) (Figure 6A). To avoid total PKA_cat_ depletion by wild-type ARHGAP36 in these studies, we also limited the cells to a 4-hour incubation in retroviral medium and a subsequent 20-hour growth phase. The fibroblasts were then lysed, and each ARHGAP36 construct and its interacting proteins were isolated by tandem affinity purification and proteolytically digested as previously described (*Ding, et al., 2016; Hsu, et al., 2019; Kanie, et al., 2017; Li, Bin, et al., 2017; Torres, et al., 2009; Wright, et al., 2011*). The resulting peptides were sequenced and quantified using tandem mass spectrometry, and spectral counts were normalized to account for variabilities in protein size, LAP tag purification efficiency, and ARHGAP36 expression. Using this approach, we identified 566 putative interactors for wild-type and/or L317P ARHGAP36 that were observed across three biological replicates (Figure 6B, Supplementary File 2). PKA_cat_ subunits were the only canonical Hh pathway regulators detected in these pulldown experiments, and they interacted with wild-type and L317P ARHGAP36 to similar extents (Figure 6B, Figure 6 – figure supplement 1). Eleven proteins preferentially bound to wild-type ARHGAP36 by at least 5-fold, with prolyl oligopeptidase-like protein PREPL and the E3 ubiquitin ligase PRAJA2 exhibiting the greatest selectivity (52- and 35-fold, respectively). PREPL displays high sequence homology to a family of serine hydrolases, though its physiological substrates and functions remain largely unknown (*Jaeken, et al., 2006; Radhakrishnan, et al., 2013; Szeltner, et al., 2005*). PRAJA2 has been shown to increase PKA activity by promoting the ubiquitination and degradation of regulatory PKA (PKA_reg_) subunits, a function that is enhanced by PKA_cat_ phosphorylation as part of a positive-feedback mechanism (*Lignitto, et al., 2011*).

**Figure 6.**
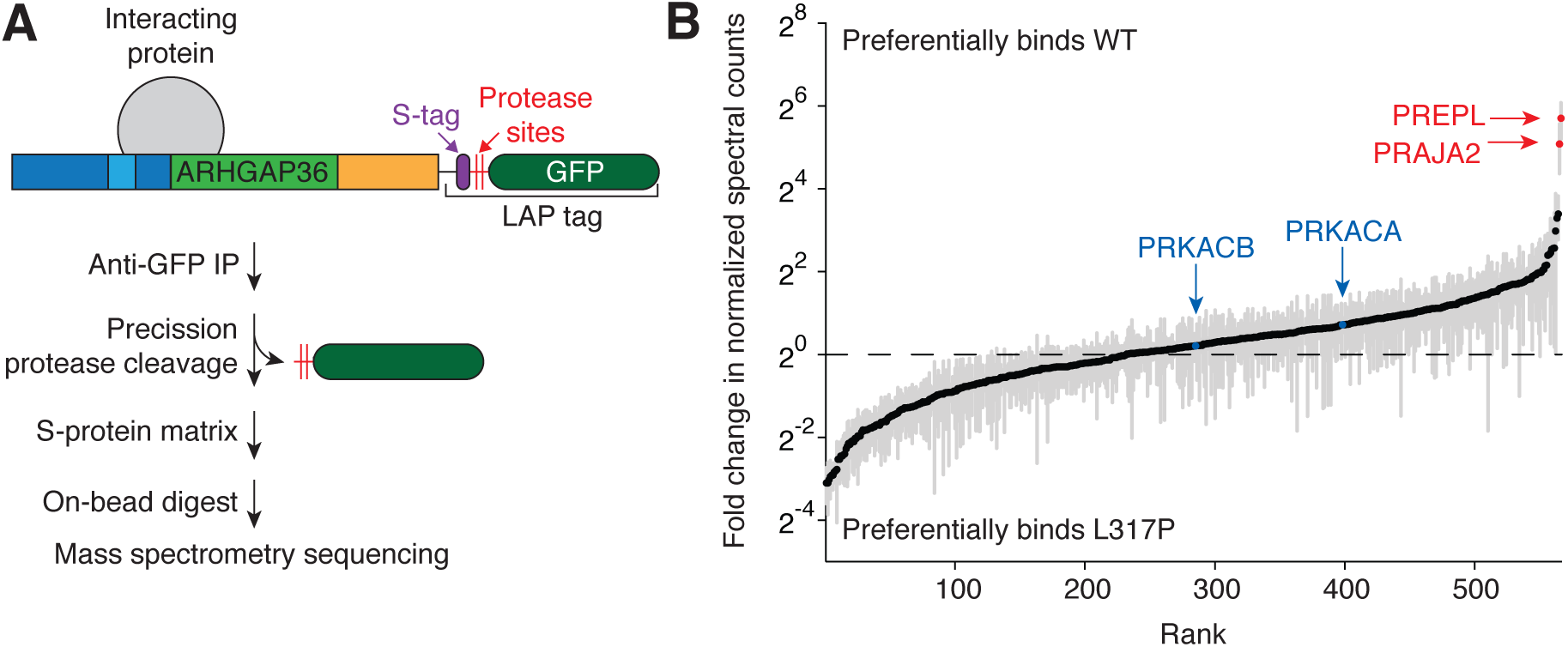
PREPL and PRAJA2 interact with ARHGAP36 in a GAP-like domain-dependent manner. (A) Schematic representation of the tandem affinity purification workflow for identifying ARHGAP36-binding proteins. (B) ARHGAP36 interactors ranked in order of their relative binding to the wild-type versus L317P proteins. Fold changes in normalized spectral counts represent the average values for three biological replicates ± s.e.m., shown as grey bars.

Both PREPL and PRAJA2 have been classified as putative ARHGAP36-binding partners in large-scale interactome studies (*Huttlin, et al., 2017; Müller, et al., 2020*); however, their functional significance relative to the other candidates in those lists has been unclear. Our comparative interactome analyses therefore corroborate the results of these prior investigations and suggest that ARHGAP36-dependent Gli activation involves PREPL and PRAJA2 functions.

## DISCUSSION

By systematically exploring the ARHGAP36 structure-activity landscape, we have identified key functional elements throughout this multidomain protein. Previous investigations identified an N-terminal arginine-rich region that is necessary and sufficient for PKA_cat_ degradation (*Eccles, et al., 2016; Rack, et al., 2014*), establishing ARHGAP36 as a novel antagonist of PKA signaling. Isoform-specific differences have also implicated N-terminal sequences in ARHGAP36 trafficking (*Rack, et al., 2014*). Our studies provide further evidence for these mechanisms of ARHGAP36 action, uncover a new functional module within the N-terminal domain, and demonstrate important regulatory roles for the GAP-like and C-terminal domains (Figure 7).

**Figure 7.**
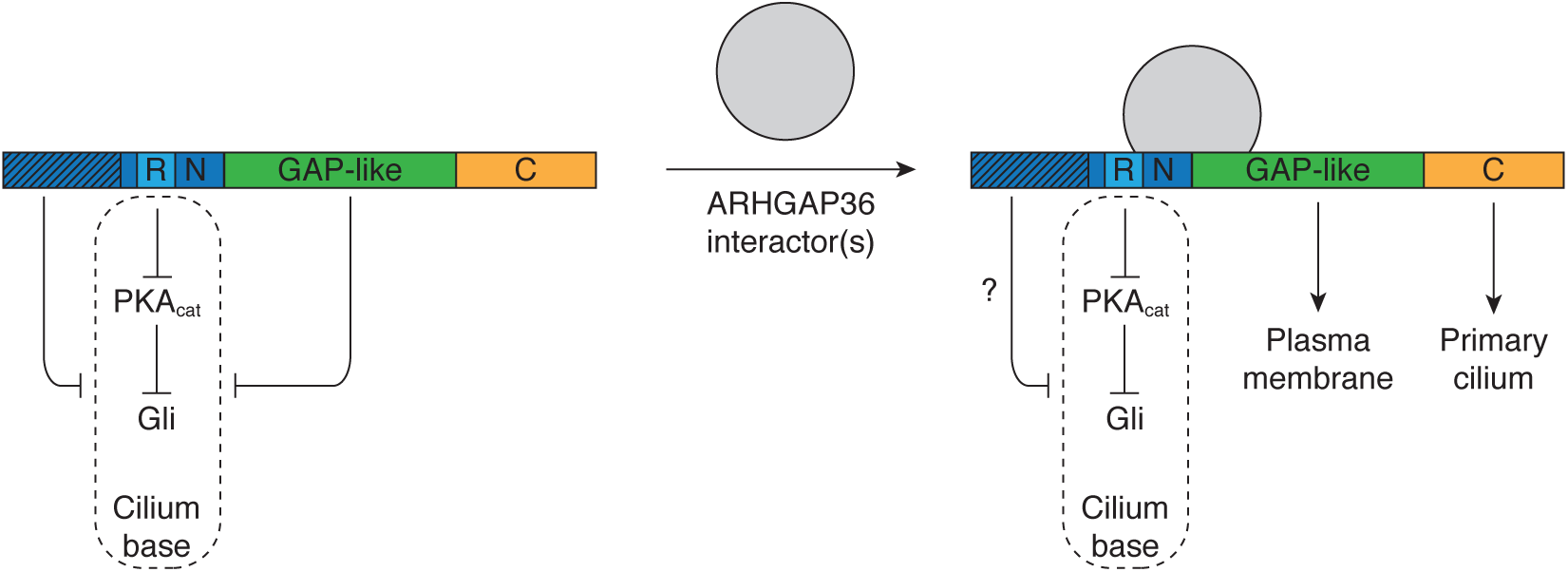
A regulatory model for ARHGAP36 function. Schematic representation of the structural elements that regulate ARHGAP36 localization and its ability to target Gli-regulating pools of PKAcat in the primary cilium. N-terminal regulatory regions that vary between ARHGAP36 isoforms are depicted with hash marks.

Among our key findings is the discovery of an N-terminal autoinhibitory region that is present in isoforms 1 and 2. In the context of isoform 2, this sequence (N2_1-105_) represses Gli1 activation by the N-terminal domain and impedes its recruitment to the plasma membrane. N2_1-105_ likely regulates these activities through distinct mechanisms, since the L86P mutation in this region diminishes the ability of isoform 2 to modulate PKA_cat_/Gli signaling without disrupting membrane localization. ARHGAP36 isoform 3, which lacks the region corresponding to N2_1-105_, not only traffics to the plasma membrane but also accumulates in the primary cilium. The ciliary localization of isoform 3 coincides with its ability to selectively degrade Golgi-localized PKA_cat_, likely due to vesicular transport between the cilium base and the Golgi (*Pedersen, et al., 2016*). Our results suggest that the Gli-regulating pool of PKA_cat_ traffics through these organelles, corroborating previous reports that Hh signaling regulates PKA_cat_ activity in the basal body (*Barzi, et al., 2009; Tuson, et al., 2011; Zhang, et al., 2019*). This mechanism of ARHGAP36-mediated Gli activation parallels the actions of other Hh pathway regulators that modulate ciliary PKA activity, such as adenylate cyclases 5 and 6 and the G-protein coupled receptor GPR161 (*Chávez, et al., 2015; Garcia-Gonzalo, et al., 2015; Moore, et al., 2016; Mukhopadhyay, et al., 2013; Vuolo, et al., 2015*). ARHGAP36 might concurrently modulate Gli function via PKA-independent mechanisms, since forskolin can only inhibit ARHGAP36-induced Gli activity by 50%, even when maximal compound doses and different levels of Gli activation are employed.

Although the ARHGAP36 N-terminal domain alone can target PKA_cat_ for degradation and harbors regulatory elements, its function is further modulated by other structures in the full-length protein. The C-terminal domain strongly counteracts the autoinhibitory activity of N2_1-105_, as does the GAP-like region to a more moderate extent. These regulatory actions likely involve protein trafficking since the GAP-like and C-terminal domains promote ARHGAP36 recruitment to the plasma membrane and primary cilium, respectively. However, we cannot rule out the possibility that the two domains influence N2_1-105_ function through additional mechanisms. Interestingly, the ability of point mutations in the GAP-like domain to abrogate Gli activation by isoforms 2 and 3 contrasts the sufficiency of N2 and N3 for this process. The mutations also render these ARHGAP36 proteins cytosolic. These differences could be explained if the unbound GAP-like domain can repress the N-terminal region conserved between N2 and N3, which contains the PKA_cat_-targeting arginine-rich motif and a plasma membrane-targeting sequence. Recruitment of specific cellular factors to the GAP-like domain could then modulate this interaction and possibly mediate other ARHGAP36 activities (see Figure 7).

Which interacting proteins might regulate or transduce ARHGAP36 function remains to be determined, but our investigations provide valuable leads. By comparing the interactomes of wild-type ARHGAP36 and an inactive mutant, we have identified several proteins that bind to ARHGAP36 in a GAP-like domain-dependent manner. Although previous studies have shown that ARHGAP36 can co-immunoprecipitate with PTCH1 (*Zhang, et al., 2019*) and SUFU (*Rack, et al., 2014*) when they are overexpressed in cells, we do not observe analogous interactions with the endogenous Hh signaling proteins. Rho GTPases were also notably absent from the ARHGAP36 pulldowns, suggesting that ARHGAP36 interacts transiently with these signaling proteins or prefers other binding partners. In contrast, PKA_cat_ associated with both wild-type and L317P ARHGAP36 with comparable efficacies, indicating that the GAP-like domain mutation abrogates PKA_cat_ degradation without compromising its binding. Among the ARHGAP36 interactors discovered in our studies, PREPL and PRAJA2 emerged as the two most sensitive to the L317P mutation in the GAP-like domain. Both proteins were candidates in previous ARHGAP36 interactome datasets (*Huttlin, et al., 2017; Müller, et al., 2020*), but the functional relevance of these factors and other ARHGAP36-binding proteins has yet to be determined. By comparing the wild-type and mutant ARHGAP36 interactomes, we can provide a functional context for these binding proteins, implicating PREPL and PRAJA2 in ARHGAP36-dependent Gli activation.

In principle, the GAP-like domain could bind directly to PREPL or PRAJA2, or it could allosterically regulate their interactions with other regions in the ARHGAP36 protein. PREPL is highly homologous to the prolyl oligopeptidase (PREP) family of serine hydrolases, but its hydrolytic targets remain uncharacterized (*Jaeken, et al., 2006; Szeltner, et al., 2005*). Putative peptidic substrates could include factors that regulate PKA_cat_ or Gli activity. Alternatively, there is evidence that PREP enzymes can modulate the metabolism of phosphoinositides (*Schulz, et al., 2002; Williams, et al., 1999*), a family of lipids with reported roles in the formation of Golgi-derived vesicles (*Heldwein, et al., 2004; Wang, Ying Jie, et al., 2003*) and the ciliary trafficking of Hh pathway regulators (*Chávez, et al., 2015; Garcia-Gonzalo, et al., 2015*). The interaction of ARHGAP36 with PRAJA2 is seemingly paradoxical since PRAJA2 has well-established roles in promoting PKA_reg_ degradation and PKA_cat_ activation (*Lignitto, et al., 2013; Lignitto, et al., 2011; Sepe, et al., 2014*). However, one possibility is that PRAJA2-mediated PKA_reg_ degradation leads to the mislocalization of Gli-regulating pools of PKA_cat_. We note that neither PREPL nor PRAJA2 were identified as hits in CRISPR knockout screens for Hh pathway regulators (*Breslow, et al., 2018; Pusapati, et al., 2018*), suggesting that they participate in an ARHGAP36-specific pathway for Gli activation. Determining how these factors contribute to ARHGAP36 action could uncover novel aspects of Gli regulation.

Taken together, our findings provide new insights into the mechanisms that underlie non-canonical Gli activation by ARHGAP36, and they provide a general framework for understanding ARHGAP36 function. In particular, our studies reveal how ARHGAP36 can translate multiple cellular inputs into distinct signaling outputs. By controlling the expression of ARHGAP36 isoforms and/or ARHGAP36-interacting proteins, cells can direct this signaling protein to specific PKA_cat_ populations and elicit tissue-specific responses. We anticipate that these mechanisms not only contribute to Gli-dependent spinal cord development and medulloblastoma progression but also other PKA_cat_-dependent processes in normal physiology and disease. Moreover, the experimental methods utilized for our study could be redeployed to elucidate these unique ARHGAP36 functions.

## METHODS

### Reagents and cell lines

Antibody sources and working dilutions are listed in Supplementary File 3. Forskolin was purchased from Calbiochem, and SAG was purchased from Tocris. SHH-LIGHT2 (*Taipale, J, et al., 2000*) and SHH-EGFP cells (*Hyman, et al., 2009*) were described previously, and NIH-3T3 and HEK-293T cells were purchased from the American Type Culture Collection.

### Expression vectors

The pDONR223 vector was provided by J. Hartley and D. Esposito. The following constructs have been described previously (*Rack, et al., 2014; Wright, et al., 2011*): Gateway cloning destination vectors pBMN-3xFLAG-IRES-mCherry-DEST, pBMN-mCherry-DEST, and pG-LAP7-DEST, Gateway cloning entry vectors pDONR223-ARHGAP36 (isoforms 1 - 5), and expression vectors pcDNA3.2-ARHGAP36 (isoform 2)-V5, pBMN-3xFLAG-IRES-mCherry, and pBMN-mCherry. pCL-ECO retrovirus packaging vector was purchased from Imegenex.

Retroviral expression vectors for FLAG-tagged ARHGAP36 isoforms 1 - 5 were produced by transferring cDNAs from the appropriate pDONR223 entry vectors into pBMN-3xFLAG-IRES-mCherry-DEST in an LR Clonase II (Invitrogen, Waltham, MA)-mediated recombination reaction. Retroviral expression vectors for mCherry-tagged ARHGAP36 isoforms 1 and 2 were produced in an analogous manner with pBMN-mCherry-DEST. Restriction sites were then inserted upstream (XhoI) and downstream (SacII) of the *ARHGAP36* sequence in the initial pBMN-ARHGAP36 (isoform 1)-mCherry and pBMN-ARHGAP36 (isoform 2)-mCherry products. This was achieved by first amplifying *ARHGAP36* cDNA from the pBMN-derived vectors (Supplementary File 3; primers 1 - 3) and then inserting the resulting PCR product into BamHI-digested pBMN-mCherry using Gibson assembly (New England Biolabs, Ipswich, MA). The resulting pBMN-ARHGAP36-mCherry vectors with XhoI and SacII restriction sites were subsequently used in all experiments described herein.

Retroviral expression vectors for ARHGAP36 isoform 2 truncation mutants were generated by amplifying the cDNA for each variant from pcDNA3.2-ARHGAP36 (isoform 2)-V5, using primers containing attB adapter sequences (Supplementary File 3; primers 4-8). The PCR products were transferred into pDONR223 in a BP Clonase II (Invitrogen)-mediated recombination reaction, and the *ARHGAP36*-derived cDNAs in these pDONR223 entry vectors were then transferred to pBMN-3xFLAG-IRES-mCherry-DEST using LR Clonase II. The resulting constructs were also used as templates to amplify cDNAs for the analogous FLAG-tagged ARHGAP36 isoform 3 truncation mutants and the downstream IRES-mCherry sequence (Supplementary File 3; primers 9 and 10). These PCR products were then inserted into XcmI-digested pBMN-ARHGAP36 (isoform 2)-3xFLAG-IRES-mCherry using Gibson assembly.

Individual ARHGAP36 isoform 2 point mutants, with the exception of L171P, were generated using site-directed mutagenesis with PfuUltra II Fusion polymerase (Agilent, Santa Clara, CA) (Supplementary File 3; primers 11-42) and either pBMN-ARHGAP36 (isoform 2)- 3xFLAG-IRES-mCherry or the pBMN-ARHGAP36 (isoform 2)-mCherry as template. ARHGAP36 isoform 2 L171P mutant constructs were generated by Gibson assembly using a XhoI- and SacII-digested pBMN-ARHGAP36 (isoform 2)-mCherry plasmid, inserts amplified from pBMN-ARHGAP36 (isoform 2)-mCherry (Supplementary File 3; primers 3 and 43-45), and a double-stranded oligonucleotide encoding the L171P mutation (Integrated DNA Technologies, Coralville, IA) (Supplementary File 3; entry 46).

Retroviral expression vectors for mCherry-tagged wild-type, C107Y, and L212P ARHGAP36 isoform 3 were generated by amplifying the cDNAs encoding this isoform from the corresponding mutant pBMN-ARHGAP36 (isoform 2)-mCherry plasmids (Supplementary File 3; primers 3 and 47) and amplifying the mCherry tag from pBMN-ARHGAP36 (isoform2)-mCherry plasmid (Supplementary File 3; primers 10 and 48). The resulting amplicons were inserted into XcmI-digested pBMN-ARHGAP36 (isoform 2)-3xFLAG-IRES-mCherry using Gibson assembly. Retroviral expression vectors for mCherry-tagged wild-type, C212Y, and L317P ARHGAP36 N2-GAP were generated by amplifying the cDNAs encoding ARHGAP36 N2-GAP from the corresponding mutant pBMN-ARHGAP36 (isoform 2)-mCherry plasmids (Supplementary File 3; primers 45 and 49) and inserting into XhoI- and SacII-digested pBMN-ARHGAP36 (isoform 2)- mCherry using Gibson assembly.

To generate retroviral expression vectors for LAP-tagged ARHGAP36 constructs, *ARHGAP36* cDNA in the pDONR223-ARHGAP36 (isoform 2) entry vector was transferred to pG- LAP7-DEST using LR Clonase II. The cDNA encoding LAP-tagged ARHGAP36 was amplified from the resulting pG-ARHGAP36 (isoform 2)-LAP7 plasmid (Supplementary File 3; primers 10 and 50) and inserted into XcmI-digested pBMN-ARHGAP36 (isoform 2)-3xFLAG-IRES-mCherry using Gibson assembly. LAP-tagged L317P ARHGAP36 isoform 2 retroviral expression constructs were generated using an analogous Gibson assembly with *ARHGAP36* cDNA amplified from pBMN-ARHGAP36 (isoform 2 L317P)-3xFLAG-IRES-mCherry (Supplementary File 3; primers 50 and 51) and LAP tag cDNA amplified from pG-ARHGAP36 (isoform 2)-LAP7 (Supplementary File 3; primers 10 and 48).

With the exception of the constructs generated by site-directed mutagenesis described above, all PCR products were generated with Phusion polymerase (New England Biolabs). All plasmids were sequence-verified.

### Retrovirus production

HEK-293T cells were seeded into individual wells of a 6-well plate at a density of 1.0 x 10^6^ cells/well. The cells were cultured for 24 hours in HEK-293T growth medium (DMEM containing 10% fetal bovine serum, 2 mM L-glutamine, 1 mM sodium pyruvate, 100 U/mL penicillin, and 0.1 mg/mL streptomycin) and then transfected as follows. pBMN vectors containing the appropriate ARHGAP36 construct (1.33 μg) and the pCL-ECO retrovirus packaging vector (0.67 μg) were diluted in OMEM medium (75 μL), and the solution was added to OMEM (75 μL) containing 6 μL Fugene HD reagent (Promega, Madison, WI). The mixture was incubated at room temperature for 10 minutes and gently added to the growth medium on the cultured cells. After 24 hours, the medium was replaced with DMEM containing 1.8 mM L-glutamine, 4% fetal bovine serum, 6% calf serum, 1 mM sodium pyruvate, 100 U/mL penicillin, and 0.1 mg/mL streptomycin. Retrovirus-containing supernatant was then collected two times at 20-hour intervals, passed through a 0.45-μm filter, and stored at −80 °C. Large-scale retrovirus production was conducted using HEK-293T cells seeded on 10-cm plates at a density of 5.0 x 10^6^ cells/well transfected with 6.48 μg of the ARHGAP36 construct, 4 μg of pCL-ECO, and 35 μL of Fugene HD in 750 μL OMEM medium.

### Generation of ARHGAP36-expressing NIH-3T3 cell lines

NIH-3T3 cells were seeded into individual wells of 24-well or 6-well plates at a density of 7.5 x 10^4^ or 2.0 x 10^5^ cells/well, respectively. The cells were cultured for 24 hours in NIH-3T3 growth medium (DMEM containing 10% calf serum, 1 mM sodium pyruvate, 100 U/mL penicillin, and 0.1 mg/mL streptomycin) and then transduced with 4 μg/mL polybrene and retrovirus for the appropriate ARHGAP36-3xFLAG-IRES-mCherry construct to achieve a multiplicity of infection (MOI) < 0.5. After 24 hours, the medium was exchanged, and the cells were expanded for fluorescence-activated cell sorting (FACS).

For FACS, the cells were washed with PBS buffer, dissociated with TrypLE (Invitrogen) for 3-5 minutes at 37 °C, and centrifuged at 106 *g* for 5 minutes at 4 °C. Cell pellets were then resuspended in FACS buffer (PBS containing 1% calf serum), passed through a 70-μm cell strainer (BD Biosciences, San Jose, CA), and added to round-bottom FACS tubes. Cell populations with comparable mCherry fluorescence intensities were then obtained using a BD FACSAria II (532-nm laser and 600-nm longpass filter, or 561-nm laser and 610/20-nm bandpass filter) or BD Influx (561-nm laser and 610/20-nm bandpass filter) sorter.

### Quantitative reverse transcription-PCR (qRT-PCR) analyses

NIH-3T3 cell lines expressing the indicated FLAG-tagged ARHGAP36 constructs were seeded into 6-well plates at a density of 5.2 x 10^5^ cells/well and cultured in NIH-3T3 growth medium. An uninfected condition was also prepared as a negative control. After two days, fully confluent cells were treated with NIH-3T3 low-serum medium (DMEM containing 0.5% calf serum, sodium pyruvate and antibiotics) with or without 10% SHH-N-conditioned medium for 30 hours. The media was replaced with ice-cold PBS, and the cells were collected by manual scraping. Each resulting cell suspension was divided into two tubes (one each for qRT-PCR and Western blot analyses) and centrifuged at 750 *g* for 7 minutes at 4 °C.

Cell pellets were prepared for qRT-PCR analyses as follows. RNA was isolated using the Monarch Total RNA miniprep kit (New England BioLabs), and equivalent amounts of RNA were used to synthesize cDNA using the SuperScript III First-Strand Synthesis System (Invitrogen). qRT-PCR was performed on a Lightcycler 480 II (Roche, Penzberg, Germany) using the following TaqMan probes: Gli1-Mm00494645_m1, Beta-2-Microglobulin-Mm00437762_m1 (Applied Biosystems, Waltham, MA). Gene expression levels were normalized to β-2-microglobulin. For ARHGAP36 isoform and truncation mutant analyses, the normalized gene expression levels were compared to that of uninfected cells, and for point mutant analyses, compared to that of wild-type ARHGAP36. The resulting gene expression levels were averaged across three biological replicates, and *P* values were determined using either a Student’s one-tailed t-test (ARHGAP36 isoform and truncation mutant analyses) or two-tailed t-test (point mutant analyses).

### Western blot analyses

Cell pellets were resuspended in 1x Laemmli sample buffer (10% glycerol, 2% SDS, 17 mM DTT, 0.01% bromophenol blue, 60 mM Tris-HCl, pH 6.8, and protease and phosphatase inhibitors (Roche)). After incubation for 20 minutes at 4 °C, cell lysates were boiled for 10 minutes and sonicated in a water bath for 15 seconds. Equivalent amounts of total protein per lysate were loaded onto Criterion XT 4-12% Bis-Tris polyacrylamide gels (Bio-Rad, Hercules, CA), transferred onto PVDF membranes (Bio-Rad), and detected using the antibodies listed in Supplementary File 3 with either SuperSignal West Dura or SuperSignal Femto kits (Pierce, Waltham, MA) and a ChemiDoc XRS imaging system (Bio-Rad). Band intensities were quantified using ImageLab software (Bio-Rad) and normalized to KPNB1 band intensities in the corresponding sample. For each replicate, the normalized band intensity in each condition was normalized to that of uninfected cells. The resulting relative band intensities for each condition was averaged across three biological replicates, and *P* values were determined using a Student’s two-tailed t-test.

### Immunofluorescence studies

The subcellular localizations of FLAG-tagged ARHGAP36 constructs were assessed as follows. NIH-3T3 cells were seeded onto individual wells of a 6-well plate at a density of 2.0 x 10^5^ cells/well. Cells were cultured for 24 hours in NIH-3T3 growth medium, then transduced with 4 μg/mL polybrene and retrovirus for the appropriate ARHGAP36-3xFLAG-IRES-mCherry construct to achieve an MOI < 0.5. After 24 hours, cells were re-seeded at a 1:8 dilution into 24- well plates containing poly-D-lysine-coated 12-mm glass coverslips and cultured for 1-2 days in growth medium. Cells were fixed in PBS containing 4% paraformaldehyde for 10 minutes at room temperature and washed 3 times with PBS. Cells were next permeabilized with PBS containing 0.5% Triton X-100 for 5 minutes, washed 2 times with PBS, and incubated in blocking buffer (PBS containing 1% BSA and 0.1% Triton X-100) for 1 hour at room temperature. The cells were then incubated for 1 hour at room temperature with primary antibodies diluted in blocking buffer, washed 4 × 5 minutes with PBS containing 0.1% Triton X-100, incubated for 1 hour with the appropriate secondary antibodies diluted in PBS containing 0.2% Triton X-100, and washed 4 × 5 minutes with PBS. The coverslips were rinsed briefly in water and mounted onto slides using Prolong Gold Antifade reagent with DAPI (Invitrogen).

The subcellular localizations of mCherry-tagged ARHGAP36 isoform 3 and N2-GAP constructs were assessed as follows. SHH-EGFP cells were seeded into individual wells of a 24- well plate at a density of 7.5 x 10^4^ cells/well. The cells were cultured for 24 hours in SHH-EGFP growth medium (NIH-3T3 growth medium containing 150 μg/mL zeocin) for 24 hours and then transduced with 4 μg/mL polybrene and retrovirus for the appropriate ARHGAP36-mCherry construct to achieve an MOI < 0.5. After 24 hours, the cells were passaged into a new 24-well plate at a 1:1.5 dilution and cultured in growth medium for an additional 2 days to achieve 100% confluency. Confluent cells were then treated for 24 hours with SHH-EGFP low-serum medium (DMEM containing 0.5% calf serum, sodium pyruvate, zeocin, and antibiotics). Cells were next passaged at a 1:3 dilution into 24-well plates containing poly-D-lysine-coated 12-mm glass coverslips and cultured for 1 day in growth medium. Cells were then fixed, blocked, immunostained, and mounted as described above. The subcellular localizations of mCherry-tagged isoform 2 constructs were similarly assessed, with the exception that the cells were passaged onto coverslip-containing 24-well plates 24 hours after retroviral transduction then cultured for two days prior to being fixed.

The effects of ARHGAP36 isoforms on PKA_cat_ localization were assessed as follows. NIH-3T3 cell lines stably expressing FLAG-tagged ARHGAP36 isoforms 2 or 3 were seeded into 24- well plates containing poly-D-lysine-coated 12-mm glass coverslips at a density of 1.2 x 10^5^ cells/well. Cells were then cultured in NIH-3T3 growth medium for 2 days, at which time the cells were fixed in PBS containing 2% paraformaldehyde for 20 minutes at room temperature and then treated with methanol for 5 minutes at –20 °C. The fixed cells were incubated in blocking buffer for 1 hour at room temperature and incubated with primary antibodies diluted in blocking buffer overnight at 4 °C. Subsequent PBS washes, secondary antibody incubation, and mounting were conducted as described above.

Fluorescence images were obtained using either a Zeiss LSM 700 or 800 confocal microscope equipped with a 63x oil-immersion objective. Maximum-intensity Z-stack projections were created using either ZEN Black (Zeiss, Oberkochen, Germany), ZEN Blue (Zeiss), or FIJI (*Schindelin, et al., 2012*) software, fluorescence intensities were adjusted using FIJI, and images were cropped using Photoshop CC (Adobe, San Jose, CA).

### Luciferase assays

SHH-LIGHT2 cells were seeded into 10-cm plates at a density of 1.0 x 10^6^ cells/plate and cultured in SHH-LIGHT2 growth medium (NIH-3T3 growth medium containing 150 μg/mL zeocin and 500 μg/mL G418). After 24 hours of growth, cells were transduced with 4 μg/mL polybrene and retrovirus for either the appropriate ARHGAP36-3xFLAG-IRES-mCherry construct or for a 3xFLAG-IRES-mCherry construct for another 24 hours. The cells were then re-seeded into individual wells of a 96-well plate at a density of 3.5 x 10^4^ cells/well. All cells were cultured for an additional 24 hours, at which time the fully confluent cells were treated for 30 hours with varying conditions of forskolin in SHH-LIGHT2 low-serum medium (DMEM containing 0.5% calf serum, 1% sodium pyruvate, zeocin, G418, and antibiotics) with or without 10% SHH-N-conditioned medium. The cells were then lysed and luciferase activities were measured using a Dual-Luciferase Reporter Assay System (Promega) on a Veritas luminometer (Turner BioSystems, Sunnyvale, CA). At least three biological replicates were conducted for each condition.

### Mutant library generation

A library encoding ARHGAP36 isoform 2 mutants was created via error-prone PCR (epPCR) using the GeneMorph II Random Mutagenesis Kit (Agilent). To determine the optimal epPCR conditions for library generation, the mutation frequency was estimated for epPCRs consisting of 15, 20, 25, or 29 cycles. All reactions were conducted according to the manufacturer’s instructions, using 1.64 μg of the pBMN-ARHGAP36 (isoform2)-mCherry plasmid as template, 0.4 μM of each primer (Supplementary File 3; primers 45 and 52), and 4% DMSO. The product yield for each condition was estimated by resolving 10% of the reaction on a 1% EtBr-agarose gel and quantifying the band intensity of the resulting amplicon with ImageLab software (Bio-Rad). The 1.6-kb amplicon was gel-extracted using the QiaQuick Gel Extraction Kit (Qiagen, Hilden, Germany) and ligated into a XhoI- and SacII-digested pBMN-ARHGAP36 (isoform 2)-mCherry vector using Gibson assembly. To estimate the mutation frequency for each epPCR-generated library, XL-10 Gold *E. coli (*Agilent) were chemically transformed with 1:4 diluted Gibson assembly products and plated onto ampicillin-agarose plates. Forty colonies from each plate were sequenced using rolling circle amplification and Sanger sequencing (Sequetech, Mountain View, CA) (Supplementary File 3; primers 53-58). High-quality *ARHGAP36* reads were aligned to the coding sequence for wild-type ARHAGP36 isoform 2, and the number of nucleotide mutations within the coding sequence was counted for each read. This analysis yielded the distribution of mutated nucleotides across the library.

Through these pilot studies, we found that the 15-cycle epPCR conditions maximized the percentage of *ARHGAP36* variants with single-nucleotide changes. We next generated a large-scale library using the 15-cycle epPCR and Gibson assembly strategy described above. The undiluted Gibson reaction (8 μL) was electroporated into 160 μL of MegaX DH10B T1^R^ Electrocomp cells (Invitrogen). The electroporated cells were immediately transferred to 480 mL of Superior Broth containing 75 μg/mL ampicillin, and 100 μL of the culture was plated on ampicillin-agar plates to estimate the number of colony-forming units. The final library was found to contain approximately 4 x 10^5^ colony-forming units, which corresponds to an equivalent number of library elements. The liquid culture was incubated at 30 °C until it reached an OD_600_ of 1, after which plasmids were isolated using the NucleoBond Xtra Midi Plus Kit (Macherey-Nagel, Düren, Germany).

Retroviral medium harboring the ARHGAP36 mutant library was generated in the following manner. One 10-cm plate of HEK-293T cells at 90% confluency was transfected with 6.5 μg of the pBMN-ARHGAP36 (isoform 2)-mCherry mutant library and 4 μg of pCL-ECO using the FuGene HD transfection reagent (Promega). The medium was replaced after 24 hours with DMEM containing 1.8 mM L-glutamine, 4% fetal bovine serum, 6% calf serum, 1% sodium pyruvate, 100 U/mL penicillin, and 0.1 mg/mL streptomycin. Retrovirus-containing supernatant was then collected two times at 24-hour intervals, passed through a 0.45-μm filter, and stored at −80 °C.

### FACS-based screening

SHH-EGFP cells were seeded onto a 15-cm plate at a density of 1.0 x 10^6^ cells/plate and cultured in SHH-EGFP growth medium for 2 days, then transduced with the retroviral library of ARHGAP36 mutants and 4 μg/mL polybrene to achieve an MOI < 0.5. After 24 hours, the cells were expanded to 4 x 15-cm plates and cultured for an additional 2 days. SHH-EGFP cells treated with 10% SHH-N-conditioned media or transduced with retrovirus encoding wild-type ARHGAP36 isoform 2 served as positive controls for flow cytometry. For negative controls, untreated SHH-EGFP cells or cells transduced with retrovirus encoding ARHGAP36 isoform 1 were used.

To isolate mCherry+ cells by FACS, the transduced SHH-EGFP cells were washed with PBS, dissociated with TrypLE for 3-5 minutes at 37° C, and centrifuged at 750 *g* for 7 minutes at 4 °C. The resulting cell pellets were resuspended in FACS buffer (PBS containing 1% calf serum), passed through a 70-μm cell strainer, and added to round-bottom FACS tubes. Cell sorting was performed on a BD FACSAria II configured with a 561-nm laser, a 595-nm longpass filter, and a 616/23-nm bandpass filter for mCherry detection. Data was collected with FACSDiva software (BD Biosciences) and analyzed using FlowJo (FlowJo, Ashland, Oregon). 2.7 x 10^7^ cells were analyzed by FACS analysis.

This first sort produced a population of 3 x 10^6^ mCherry+ cells, which were cultured for 3 days until they reached 100% confluency. The cells were then cultured in SHH-EGFP low-serum medium to enable ARHGAP36-mediated Gli activation. After 24 hours, cells were expanded at a 1:2 dilution, cultured for 24 hours to achieve full confluency, and serum-starved again for another 24 hours.

To isolate cells expressing inactive forms of ARHGAP36 isoform 2 (mCherry+/EGFP–), 1.1 x 10^7^ cells from the expanded mCherry+ population were washed and dissociated as described above. Approximately 3 x 10^6^ cells were separated and expanded to 4 x 10^7^ cells to establish a pre-selection population, which was then washed with PBS, dissociated, and pelleted by centrifugation at 750 *g* for 7 minutes at 4 °C. The pellet was stored at –80 °C until used for genomic DNA extraction. The remaining 8 x 10^6^ mCherry+ cells were analyzed by FACS to select for those expressing inactive ARHGAP36-mCherry mutants. Cells were sorted as described above using the BD FACSAria II configured with a 488-nm dye laser, a 495-nm longpass filter, and a 530/30-nm bandpass filter for EGFP detection and the laser/filter configurations described above for mCherry detection. Approximately 4 x 10^5^ mCherry+/EGFP– cells were obtained from this second sort, and they were cultured for 2 days to reach full confluency and then subjected to 2 rounds of serum starvation. FACS sorting of this enriched population yielded 1.6 x 10^5^ mCherry+/EGFP– cells, which were expanded, frozen in SHH-EGFP growth medium containing 10% DMSO, and stored in liquid nitrogen.

We then assessed if the mCherry+/EGFP– cells were still capable of Gli-dependent EGFP expression under canonical Hh pathway activation conditions. Frozen aliquots of cells from the third sort were thawed, expanded, and subjected to two rounds of serum starvation. Approximately 1.0 x 10^7^ cells were sorted with a BD FACSAria IIu configured with a 488-nm laser, a 502-nm longpass filter, and a 525/50-nm bandpass filter for EGFP detection and a 488-nm laser, a 595-nm longpass filter, and a 610/20-nm bandpass filter for mCherry detection. 1.0 x 10^6^ mCherry+/EGFP– cells were collected and cultured for 7 days to achieve full confluency. The cells were then treated with 200 nM SAG in SHH-EGFP low-serum medium for 24 hours, expanded at a 1:2 dilution, cultured for 24 hours to enable full confluency, and treated again with 200 nM SAG for 24 hours. 2 x 10^7^ cells were then sorted with the BD FACSAria IIu to obtain 3 x 10^6^ mCherry+/EGFP+ cells. This selected population was expanded to 5 x 10^6^ cells, which were then washed with PBS, dissociated, and pelleted by centrifugation at 750 *g* for 7 minutes at 4 °C. The pellet was stored at –80 °C until used for genomic DNA extraction.

### Deep-sequencing analyses of pre- and post-selection pools

Genomic DNA was extracted from frozen pellets using the QIAamp DNA Blood Maxi Kit (Qiagen) according to manufacturer’s instructions. For each sample, *ARHGAP36* inserts were isolated from genomic DNA by PCR using 500 ng of genomic DNA, 0.5 μM each of primer (Tables S3, primers 59 and 60), 0.2 μM dNTP mix, and Phusion High-Fidelity DNA Polymerase in HF buffer (New England BioLabs). A total of 182 PCRs were used to isolate *ARHGAP36* inserts from 91 μg of pre-selection genomic DNA, while 89 PCRs were used to isolate inserts from 45 μg of post-selection genomic DNA. For each condition, the respective reactions were pooled, purified using the QIAquick PCR Purification Kit (Qiagen), and resolved on a 0.8% EtBr–agarose gel. The 1.6-kb amplicon was then gel-extracted using the QIAquick Gel Extraction Kit. Amplicons were quantified using a Bioanalyzer 2100 with high-sensitivity DNA kits (Agilent), sheared into 150-bp fragments with an S220 focused-ultrasonicator (Covaris, Woburn, MA), and sequenced on a NextSeq 500 Sequencer using High-Output v2 kits (Illumina, San Diego, CA). Raw data have been deposited into the Dryad Digital Repository with the dataset identifier 10.5061/dryad.dz08kprv9.

FASTQ files were aligned to the wild-type *ARHGAP36* coding sequence (Bowtie2 v2.3 (*Langmead, et al., 2012*) and sorted by read name (SAMtools v1.3.1) (*Li, Heng, et al., 2009*). The resulting mapped and sorted reads were then analyzed with an in-house Python script (Supplementary File 4). Briefly, high-quality reads were aligned to the wild-type *ARHGAP36* coding sequence. Reads with greater than 3 high-quality mutations or with internal stop codons were discarded from further analysis. The remaining reads were translated, identifying the ARHGAP36 amino acid mutations present in the population. For each residue, the fold-change in its mutation frequency between the pre- and post-selection populations was calculated.

### Flow cytometry-based assays

SHH-EGFP cells were seeded into individual wells of a 24-well plate at a density of 7.5 x 10^4^ cells/well and cultured for 24 hours in SHH-EGFP growth medium. The cells were then transduced with retrovirus for the appropriate ARHGAP36-mCherry construct and 4 μg/mL polybrene to achieve an MOI < 0.5. An uninfected condition was also prepared as a negative control. After 24 hours, the cells were passaged onto a new 24-well well at a 1:1.5 dilution and cultured in growth medium for an additional 2 days to achieve 100% confluency. Confluent cells were then treated with SHH-EGFP low-serum medium for 24 hours. Cells were then passaged at a 1:1.5 dilution onto a new 24-well well and cultured for 24 hours to achieve full confluency for a second round of 24-hour serum-starvation with or without SHH-N-conditioned medium.

For flow cytometry analyses, the cells were washed with PBS, dissociated with TrypLE for 3-5 minutes at 37 °C, and centrifuged 750 *g* for 7 minutes at 4 °C. Cell pellets were resuspended in FACS buffer (PBS containing 1% calf serum) and analyzed on a DxP FACScan (561-nm laser and 616/25-nm bandpass filter for mCherry detection; 488-nm laser, 560-nm shortpass filter, and 525/50-nm bandpass filter for EGFP detection) or a BD LSRII (561-nm laser, 600-nm longpass band filter, and a 610/20-nm bandpass filter for mCherry detection; 488-nm laser, 505-nm longpass band filter, and 525/50-nm bandpass filter for EGFP detection). Data was collected with Cypod (Cytek, Fremont, CA) and FACSDiva software and analyzed using FlowJo. Fluorescence data was collected for at least 2.5 x 10^4^ cells, and three biological replicates were analyzed for each condition.

Data analyses for all conditions except for the uninfected controls excluded mCherry– cells, which is indicative of a lack of ARHGAP36 expression, and the median EGFP fluorescence was calculated to measure Gli activity in each condition. For each replicate, the median EGFP fluorescence was normalized to that of wild-type ARHGAP36-expressing cells, and a Student’s one-tailed t-test was used to identify mutations that significantly altered median EGFP fluorescence levels (*P* ≤ 0.05).

### Tandem affinity purification and quantitative proteomics

NIH-3T3 cells were seeded onto 15-cm plates (6 per condition) at a density of 2 x 10^6^ cells/plate and cultured in NIH-3T3 growth medium. After 24 hours, cells were transduced with retrovirus for either wild-type or L317P ARHGAP36 with a C-terminal LAP tag and 4 μg/mL polybrene to achieve an MOI > 1.5. The media was exchanged for growth medium after 4 hours, and cells were cultured for an additional 20 hours. Cells were then washed with cold PBS, manually scraped off each dish, and transferred into Falcon tubes. Cell suspensions expressing the same ARHGAP36 construct were combined, and 0.5% of the resulting pool was reserved for downstream flow cytometry analyses to confirm the MOI. The remaining cells were centrifuged at 750 *g* for 7 minutes at 4 °C. The supernatant was aspirated, and the remaining cell pellet was flash frozen in liquid nitrogen and stored at –80 °C prior to LAP-tagged mediated tandem affinity purification. Three biological replicates were conducted for each wild-type and L317P ARHGAP36-LAP comparison.

Tandem affinity purifications and mass spectrometry analyses were conducted as described previously (*Ding, et al., 2016; Hsu, et al., 2019; Kanie, et al., 2017; Li, Bin, et al., 2017; Torres, et al., 2009; Wright, et al., 2011*). Pellets of ARHGAP36-LAP-expressing cells were re-suspended in LAP resuspension buffer (300 mM KCl, 50 mM HEPES-KOH (pH 7.4), 1 mM EGTA, 1 mM MgCl_2_, 10% glycerol, 0.5 mM DTT, and protease inhibitors (Thermo Scientific)). Cells were lysed with the gradual addition of 10% NP-40 to a final concentration of 0.3%, followed by a 10- minute incubation at 4 °C. The lysate was then centrifuged at 27,000 *g* at 4 °C for 10 minutes, and the resulting supernatant was centrifuged at 100,000 *g* for 1 hour at 4 °C. The high-speed supernatant was next incubated with anti-GFP-antibody-coupled beads (*Ding, et al., 2016; Hsu, et al., 2019; Kanie, et al., 2017; Li, Bin, et al., 2017; Torres, et al., 2009; Wright, et al., 2011*) for 1 hour at 4 °C to capture GFP-tagged proteins. The beads were washed five times with LAP200N buffer (200 mM KCl, 50 mM HEPES-KOH (pH 7.4), 1 mM EGTA, 1 mM MgCl_2_, 10% glycerol, protease inhibitors, and 0.05% NP40) and incubated with PreScission protease in LAP200N buffer at 4 °C for 16 hours. All subsequent steps were performed in a laminar flow hood. PreScission protease-eluted supernatant was added to S-protein agarose beads (EMD Millipore, Burlington, MA) and incubated rocking for 3 hours at 4 °C. S-protein agarose beads were then washed three times with LAP200N buffer and twice with LAP100 buffer (100 mM KCl, 50 mM HEPES-KOH (pH 7.4), 1 mM EGTA and 10% glycerol). Beads were stored in 50mM HEPES (pH 7.5), 1 mM EGTA, 1 mM MgCl_2_, 10% glycerol at 4 °C prior to on-bead digestion.

Proteins were eluted from S-protein agarose beads with an on-bead reduction, alkylation, and tryptic digestion as follows. Samples were reduced with 10 mM DTT in ammonium bicarbonate for an initial 5-minute incubation at 55 °C followed by 25 minutes at room temperature. The proteins were then alkylated with a 30-minute incubation in 30 mM acrylamide at room temperature, and finally eluted from the beads with an overnight digest performed at room temperature using Trypsin/LysC (Promega) and 0.02% ProteaseMax (Promega). The digests were acidified with 1% formic acid, de-salted with C18 Monospin reversed phase columns (GL Sciences, Tokyo, Japan), dried on a SpeedVac, and reconstituted in 12.5 μL of 2% acetonitrile and 0.1% formic acid. 4 μL of each sample were used for liquid-chromatography-mass spectrometry analyses performed on an Acquity M-Class UPLC (Waters Corporation, Milford, MA) and either an Orbitrap Q-Exactive HFX mass spectrometer (Thermo Scientific, San Jose, CA) or an Orbitrap Fusion Tribrid mass spectrometer (Thermo Scientific). For each biological replicate, the sample from the ARHGAP36 L317P-expressing cells was run immediately before that of the wild-type ARHGAP36-expressing cells. Analysis of the resulting .RAW data files was conducted using Byonic (Protein Metrics, San Carlos, CA), with the assumption of tryptic proteolysis and a maximum allowance of two missed cleavage sites. Precursor and MS/MS fragment mass accuracies were held within 12 ppm and 0.4 Da, respectively. A false discovery rate of 1% was used for protein identification (*Elias, et al., 2007*). Raw data have been deposited to the ProteomeXchange Consortium via the PRIDE partner repository with the dataset identifier PXD019056 and 10.6019/PXD019056.

The resulting list of identified proteins was compared to an NCBI FASTA database containing all mouse proteomic isoforms with the exception of the tandem affinity bait construct sequence and common contaminant proteins. Post-processing of spectral counts was conducted with an in-house R script (Supplementary File 5). For each protein, spectral counts detected across all isoforms were combined and normalized to the mean amino acid length of all isoforms. The resulting normalized spectral count was divided by the sum of normalized spectral counts calculated for all proteins in the pulldown sample, generating a normalized spectral abundance factor (NSAF) for each protein. To account for variability in bait ARHGAP36 expression across different pulldown samples, the NSAF of each protein was divided by that of the bait ARHGAP36 (relative NSAF). Proteins that were detected in all biological replicates for a given condition were tabulated, resulting in a dataset of 566 candidate ARHGAP36-binding proteins. For each protein, the fold-change in relative NSAF between the wild-type and L317P mutant ARHGAP36 pulldown samples of the same replicate was calculated. The average fold-change in relative NSAF across of all three replicates was then calculated for each protein.

To assess the robustness of our comparative interactome analyses, we calculated a modified Z score that compares the protein enrichment in either interactome (wild-type vs. L317P ARHGAP36) against the experimental variability in protein abundance measurements. Protein enrichment was represented by the log_2_-transformation of the average fold change in relative NSAF between wild-type and L317P ARHGAP36 interactomes [log_2_ (WT NSAF:L317 NSAF)]. We estimated the error in measuring protein abundance in a given interactome by normalizing the relative NSAFs for each replicate to the average value across all three replicates (mean-normalized relative NSAFs). This transformation approximates how much the variation between replicates can contribute to an apparent fold change. To place equal weight on upward and downward variations from the mean, we calculated the absolute value of the log_2_-transformed mean-normalized relative NSAFs. These calculations were conducted for the relative NSAFs of a protein in both the wild-type and L317P mutant ARHGAP36 interactomes, and the two resulting values were summed to produce a final error estimate. The modified Z score of each protein was calculated by dividing the log_2_ (WT NSAF:L317 NSAF) by the final error estimate and then calculating the absolute value of the resulting quotient. Proteins with higher modified Z scores are those that are enriched in a given interactome to a degree that is greater than the estimated experimental error.

### Statistical analyses

Biological replicates are defined as experimental samples that are capable of biological variance, and technical replicates are defined as those for which experimental variance is solely dependent on measurement accuracy.

## SUPPLEMENTAL DATA

Supplementary File 1. Mutation frequencies in pre- and post-selection populations

Supplementary File 2. Wild-type and L317P ARHGAP36 isoform 2 interactomes

Supplementary File 3. Antibody and primer resources

Supplementary File 4. Python script for ARHGAP36 mutagenesis screen analysis

Supplementary File 5. R script for ARHGAP36-LAP mass spectrometry analysis

## ACKNOWLEDGEMENTS

This work was supported by the National Institutes of Health (R35 GM127030 to J.K.C.; S10 RR025518-01 and S10 RR027431-01 to the Stanford Shared FACS Facility; and P30 CA124435 to Stanford University Mass Spectrometry), the Rachel Molly Markoff Foundation (J.K.C.), and an Alex’s Lemonade Stand Foundation Young Investigator Award (J.N.). Cell sorting and flow cytometry analyses were conducted at the Stanford Shared FACS Facility. Deep sequencing analyses were performed at the Stanford Functional Genomics Facility. Quantitative proteomics were executed at the Vincent Coates Foundation Mass Spectrometry Laboratory, Stanford University Mass Spectrometry.

## AUTHOR CONTRIBUTIONS

P.R. N. designed and conducted experiments, analyzed the data, and wrote the paper. T.K. developed software to analyze deep sequencing data. N.A.M. performed tandem affinity purifications. J.N. generated expression vectors for ARHGAP36 isoforms and isoform 2 truncation mutants. J.D. contributed to the development of software to analyze proteomic data. P.K.J. provided resources and supervision for proteomics experiments. J.K.C. designed the experiments, analyzed the data, and wrote the paper.

---

**Figure 4 – figure supplement 1.**
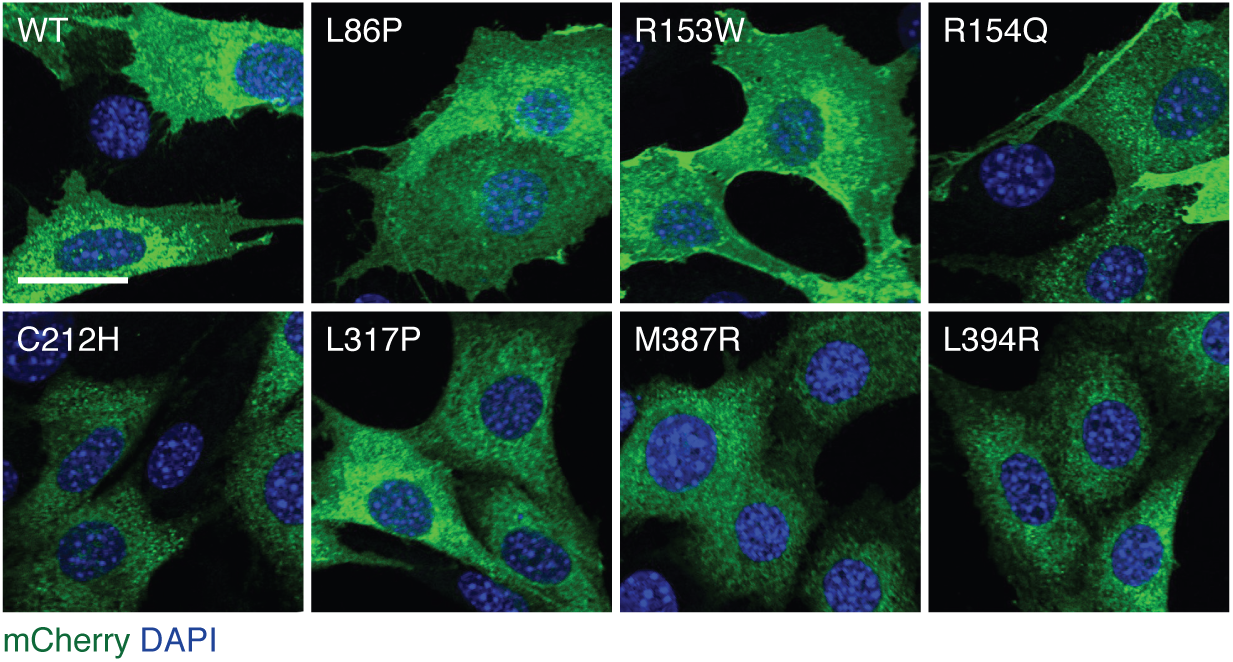
Subcellular localization of ARHGAP36 isoform 2 point mutants. Representative immunofluorescence micrographs of SHH-EGFP cells retrovirally transduced with the indicated mCherry-tagged ARHGAP36 constructs. Scale bar: 20 μm.

**Figure 6 – figure supplement 1.**
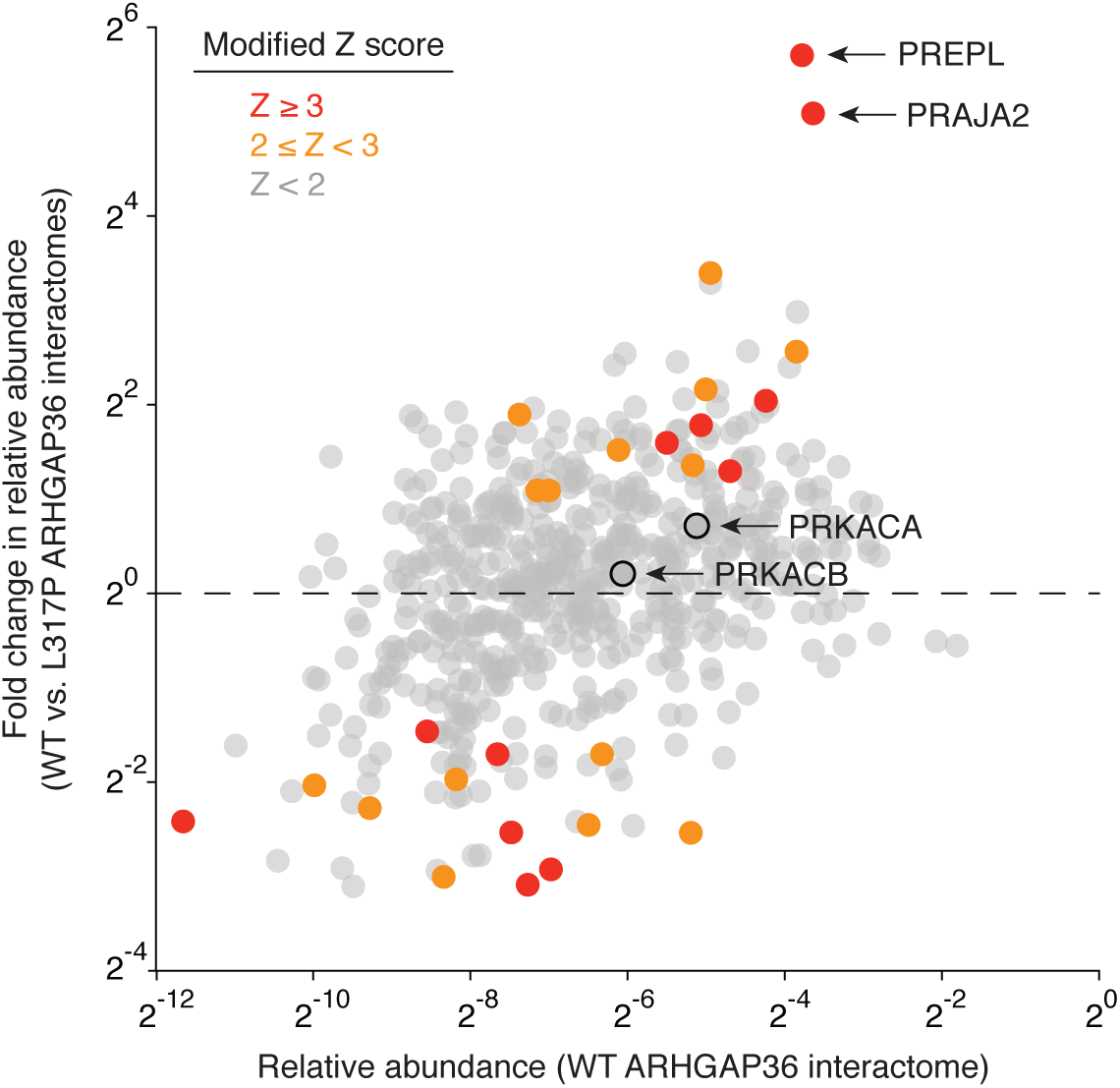
Comparative analyses of the wild-type and L317P ARHGAP36 interactomes. Scatter plot of ARHGAP36 isoform 2-binding proteins according to their normalized spectral abundance in the wild-type and L317P interactomes. Each data point represents the average value for three biological replicates, and a modified Z-score was calculated for each fold change in abundance (see Methods).

## REFERENCES

Ågren, M., Kogerman, P., Kleman, M.I., Wessling, M. & Toftgård, R. 2004. Expression of the PTCH1 tumor suppressor gene is regulated by alternative promoters and a single functional Gli-binding site. Gene 330: 101–114. doi: https://doi.org/10.1016/j.gene.2004.01.010.

Amin, E. et al. 2016. Deciphering the molecular and functional basis of RHOGAP family proteins: A systematic approach toward selective inactivation of RHO family proteins. Journal of Biological Chemistry 291: 20353–20371. doi: https://doi.org/10.1074/jbc.M116.736967.

Bai, C.B., Stephen, D. & Joyner, A.L. 2004. All mouse ventral spinal cord patterning by Hedgehog is Gli dependent and involves an activator function of Gli3. Developmental Cell 6: 103–115. doi: https://doi.org/10.1016/S1534-5807(03)00394-0.

Barzi, M., Berenguer, J., Menendez, A., Alvarez-Rodriguez, R. & Pons, S. 2009. Sonic-hedgehog-mediated proliferation requires the localization of PKA to the cilium base. Journal of Cell Science 123: 62–69. doi: https://doi.org/10.1242/jcs.060020, PMID: 5661997.

Beauchamp, E. et al. 2009. GLI1 is a direct transcriptional target of EWS-FLI1 oncoprotein. The Journal of Biological Chemistry 284: 9074–9082. doi: https://doi.org/10.1074/jbc.M806233200, PMID: 19189974.

Beckmann, P.J. et al. 2019. Sleeping beauty insertional mutagenesis reveals important genetic drivers of central nervous system embryonal tumors. Cancer Research 79: 905–917. doi: https://doi.org/10.1158/0008-5472.CAN-18-1261, PMID: 30674530.

Breslow, D.K. et al. 2018. A CRISPR-based screen for Hedgehog signaling provides insights into ciliary function and ciliopathies. Nature Genetics 50: 460–471. doi: https://doi.org/10.1038/s41588-018-0054-7.

Briscoe, J., Pierani, A., Jessell, T.M. & Ericson, J. 2000. A homeodomain protein code specifies progenitor cell identity and neuronal fate in the ventral neural tube. Cell 101: 435–445. doi: https://doi.org/10.1016/S0092-8674(00)80853-3, PMID: 10830170.

Buonamici, S. et al. 2010. Interfering with resistance to smoothened antagonists by inhibition of the PI3K pathway in medulloblastoma. Sci. Transl. Med. 10.1126/scitranslmed.3001599 51-70. doi: https://doi.org/10.1126/scitranslmed.3001599, PMID: 20881279.

Chávez, M. et al. 2015. Modulation of ciliary phosphoinositide content regulates trafficking and Sonic Hedgehog signaling output. Developmental Cell 34: 338–350. doi: https://doi.org/10.1016/j.devcel.2015.06.016.

Chen, J.K., Taipale, J., Young, K.E., Maiti, T. & Beachy, P.A. 2002. Small molecule modulation of Smoothened activity. Proceedings of the National Academy of Sciences of the United States of America 99: 14071–14076. doi: https://doi.org/10.1073/pnas.182542899.

Dai, P. et al. 1999. Sonic hedgehog-induced activation of the Gli1 promoter is mediated by GLI3. Journal of Biological Chemistry 274: 8143–8152. doi: https://doi.org/10.1074/jbc.274.12.8143.

Dennler, S. et al. 2007. Induction of sonic hedgehog mediators by transforming growth factor-beta: Smad3-dependent activation of Gli2 and Gli1 expression in vitro and in vivo. Cancer Research 67: 6981–6986. doi: https://doi.org/10.1158/0008-5472.CAN-07-0491, PMID: 17638910.

Ding, S. et al. 2016. Comparative proteomics reveals strain-specific β-TrCP degradation via rotavirus NSP1 hijacking a host Cullin-3-Rbx1 complex. PLoS Pathogens 12: e1005929. doi: https://doi.org/10.1371/journal.ppat.1005929.

Dorn, K.V., Hughes, C.E. & Rohatgi, R. 2012. A Smoothened-Evc2 complex transduces the Hedgehog signal at primary cilia. Developmental Cell 23: 823–835. doi: https://doi.org/10.1016/j.devcel.2012.07.004.

Eccles, R.L. et al. 2016. Bimodal antagonism of PKA signalling by ARHGAP36. Nature Communications 7: 12963. doi: https://doi.org/10.1038/ncomms12963, PMID: 27713425.

Elias, J.E. & Gygi, S.P. 2007. Target-decoy search strategy for increased confidence in large-scale protein identifications by mass spectrometry. Nature Methods 4: 207–214. doi: https://doi.org/10.1038/nmeth1019.

Elsawa, S.F. et al. 2011. GLI2 transcription factor mediates cytokine cross-talk in the tumor microenvironment. Journal of Biological Chemistry 286: 21524–21534. doi: https://doi.org/10.1074/jbc.M111.234146.

Faucherre, A. et al. 2003. Lowe syndrome protein OCRL1 interacts with Rac GTPase in the trans-Golgi network. Human Molecular Genetics 12: 2449–2456. doi: https://doi.org/10.1093/hmg/ddg250.

Flora, A., Klisch, T.J., Schuster, G. & Zoghbi, H.Y. 2009. Deletion of Atoh1 disrupts Sonic Hedgehog signaling in the developing cerebellum and prevents medulloblastoma. Science 326: 1424–1427. doi: https://doi.org/10.1126/science.1181453, PMID: 19965762.

Garcia-Gonzalo, F.R. et al. 2015. Phosphoinositides regulate ciliary protein trafficking to modulate Hedgehog signaling. Developmental Cell 34: 400–409. doi: https://doi.org/10.1016/j.devcel.2015.08.001.

Han, B. et al. 2015. FOXC1 activates Smoothened-independent Hedgehog signaling in basal-like breast cancer. Cell Reports 13: 1046–1058. doi: https://doi.org/10.1016/j.celrep.2015.09.063.

Haycraft, C.J. et al. 2005. Gli2 and Gli3 localize to cilia and require the intraflagellar transport protein Polaris for processing and function. PLoS Genetics 1: e53. doi: https://doi.org/10.1371/journal.pgen.0010053.

Heldwein, E.E. et al. 2004. Crystal structure of the clathrin adaptor protein 1 core. Proceedings of the National Academy of Sciences of the United States of America 101: 14108–14113. doi: https://doi.org/10.1073/pnas.0406102101.

Hill, P., Götz, K. & Rüther, U. 2009. A SHH-independent regulation of Gli3 is a significant determinant of anteroposterior patterning of the limb bud. Developmental Biology 328: 506–516. doi: https://doi.org/10.1016/j.ydbio.2009.02.017.

Hsu, J. et al. 2019. E2F4 regulates transcriptional activation in mouse embryonic stem cells independently of the RB family. Nature Communications 10: 2939. doi: https://doi.org/10.1038/s41467-019-10901-x.

Huangfu, D. & Anderson, K.V. 2005. Cilia and Hedgehog responsiveness in the mouse. Proceedings of the National Academy of Sciences of the United States of America 102: 11325–11330. doi: https://doi.org/10.1073/pnas.0505328102.

Hui, C.-c. & Angers, S. 2011. Gli proteins in development and disease. Annual Review of Cell and Developmental Biology 27: 513–537. doi: https://doi.org/10.1146/annurev-cellbio-092910-154048, PMID: 21801010.

Humke, E.W., Dorn, K.V., Milenkovic, L., Scott, M.P. & Rohatgi, R. 2010. The output of Hedgehog signaling is controlled by the dynamic association between Suppressor of Fused and the Gli proteins. Genes & Development 24: 670–682. doi: https://doi.org/10.1101/gad.1902910, PMID: 20360384.

Huttlin, E.L. et al. 2017. Architecture of the human interactome defines protein communities and disease networks. Nature 545: 505–509. doi: https://doi.org/10.1038/nature22366, PMID: 28514442.

Hyman, J.M. et al. 2009. Small-molecule inhibitors reveal multiple strategies for Hedgehog pathway blockade. Proceedings of the National Academy of Sciences of the United States of America 106: 14132–14137. doi: https://doi.org/10.1073/pnas.0907134106, PMID: 19666565.

Incardona, J.P. et al. 2000. Receptor-mediated endocytosis of soluble and membrane-tethered sonic hedgehog by patched-1. Proceedings of the National Academy of Sciences of the United States of America 97: 12044–12049. doi: https://doi.org/10.1073/pnas.220251997.

Jaeken, J. et al. 2006. Deletion of PREPL, a gene encoding a putative serine oligopeptidase, in patients with hypotonia-cystinuria syndrome. American Journal of Human Genetics 78: 38–51. doi: https://doi.org/10.1086/498852, PMID: 16385448.

Kanie, T. et al. 2017. The CEP19-RABL2 GTPase complex binds IFT-B to Iinitiate intraflagellar transport at the ciliary base. Developmental Cell 42: 22–36.e12. doi: https://doi.org/10.1016/j.devcel.2017.05.016.

Kasper, M. et al. 2006. Selective modulation of Hedgehog/GLI target gene expression by epidermal growth factor signaling in human keratinocytes. Molecular and Cellular Biology 26: 6283–6298. doi: https://doi.org/10.1128/mcb.02317-05.

Kim, J., Kato, M. & Beachy, P.A. 2009. Gli2 trafficking links Hedgehog-dependent activation of Smoothened in the primary cilium to transcriptional activation in the nucleus. Proceedings of the National Academy of Sciences of the United States of America 106: 21666–21671. doi: https://doi.org/10.1073/pnas.0912180106, PMID: 19996169.

Langmead, B. & Salzberg, S.L. 2012. Fast gapped-read alignment with Bowtie 2. Nature Methods 9: 357–359. doi: https://doi.org/10.1038/nmeth.1923, PMID: 22388286.

Lee, Y.s. et al. 2019. A computational framework for genome-wide characterization of the human disease landscape. Cell Systems 8: 152–162. doi: https://doi.org/10.1016/j.cels.2018.12.010.

Lewis, P.M., Gritli-Linde, A., Smeyne, R., Kottmann, A. & McMahon, A.P. 2004. Sonic hedgehog signaling is required for expansion of granule neuron precursors and patterning of the mouse cerebellum. Developmental Biology 270: 393–410. doi: https://doi.org/10.1016/j.ydbio.2004.03.007, PMID: 15183722.

Li, B. et al. 2017. Drebrin restricts rotavirus entry by inhibiting dynamin-mediated endocytosis. Proceedings of the National Academy of Sciences of the United States of America 114: E3642–E3651. doi: https://doi.org/10.1073/pnas.1619266114.

Li, H. et al. 2009. The Sequence Alignment/Map format and SAMtools. Bioinformatics 25: 2078- 2079. doi: https://doi.org/10.1093/bioinformatics/btp352, PMID: 19505943.

Lignitto, L. et al. 2013. Proteolysis of MOB1 by the ubiquitin ligase praja2 attenuates Hippo signalling and supports glioblastoma growth. Nature Communications 4: 1822. doi: https://doi.org/10.1038/ncomms2791.

Lignitto, L. et al. 2011. Control of PKA stability and signalling by the RING ligase praja2. Nature Cell Biology 13: 412–422. doi: https://doi.org/10.1038/ncb2209.

Liu, A., Wang, B. & Niswander, L.A. 2005. Mouse intraflagellar transport proteins regulate both the activator and repressor functions of Gli transcription factors. Development 132: 3103–3111. doi: https://doi.org/10.1242/dev.01894.

Liu, Z., Li, T., Reinhold, M.I. & Naski, M.C. 2014. MEK1-RSK2 contributes to Hedgehog signaling by stabilizing GLI2 transcription factor and inhibiting ubiquitination. Oncogene 33: 65–73. doi: https://doi.org/10.1038/onc.2012.544, PMID: 23208494.

Long, J. et al. 2014. The BET bromodomain inhibitor I-BET151 acts downstream of smoothened protein to abrogate the growth of hedgehog protein-driven cancers. Journal of Biological Chemistry 289: 35494–35502. doi: https://doi.org/10.1074/jbc.M114.595348.

Marchesi, S. et al. 2014. DEPDC1B coordinates de-adhesion events and cell-cycle progression at mitosis. Developmental Cell 31: 420–433. doi: https://doi.org/10.1016/j.devcel.2014.09.009.

May, S.R. et al. 2005. Loss of the retrograde motor for IFT disrupts localization of Smo to cilia and prevents the expression of both activator and repressor functions of Gli. Developmental Biology 287: 378–389. doi: https://doi.org/10.1016/j.ydbio.2005.08.050, PMID: 16229832.

Moon, S. 2003. Rho GTPase-activating proteins in cell regulation. Trends in Cell Biology 13: 13–22. doi: https://doi.org/10.1016/S0962-8924(02)00004-1.

Moore, B.S. et al. 2016. Cilia have high cAMP levels that are inhibited by Sonic Hedgehog-regulated calcium dynamics. Proceedings of the National Academy of Sciences of the United States of America 113: 13069–13074. doi: https://doi.org/10.1073/pnas.1602393113, PMID: 27799542.

Mukhopadhyay, S. et al. 2013. The ciliary G-protein-coupled receptor Gpr161 negatively regulates the Sonic hedgehog pathway via cAMP signaling. Cell 152: 210–223. doi: https://doi.org/10.1016/j.cell.2012.12.026, PMID: 23332756.

Müller, P.M. et al. 2020. Systems analysis of RhoGEF and RhoGAP regulatory proteins reveals spatially organized RAC1 signalling from integrin adhesions. Nature Cell Biology 22: 498– 511. doi: https://doi.org/10.1038/s41556-020-0488-x, PMID: 32203420.

Murone, M., Rosenthal, A. & De Sauvage, F.J. 1999. Sonic hedgehog signaling by the patched-smoothened receptor complex. Current Biology 9: 76–84. doi: https://doi.org/10.1016/S0960-9822(99)80018-9, PMID: 10021362.

Nam, H. et al. 2019. Critical roles of ARHGAP36 as a signal transduction mediator of shh pathway in lateral motor columnar specification. eLife 8: e46683. doi: https://doi.org/10.7554/eLife.46683.

Pan, Y., Bai, C.B., Joyner, A.L. & Wang, B. 2006. Sonic hedgehog signaling regulates Gli2 transcriptional activity by suppressing its processing and degradation. Molecular and Cellular Biology 26: 3365–3377. doi: https://doi.org/10.1128/MCB.26.9.3365-3377.2006, PMID: 16611981.

Pan, Y. & Wang, B. 2007. A novel protein-processing domain in Gli2 and Gli3 differentially blocks complete protein degradation by the proteasome. The Journal of Biological Chemistry 282: 10846–10852. doi: https://doi.org/10.1074/jbc.M608599200, PMID: 17283082.

Pedersen, L.B., Mogensen, J.B. & Christensen, S.T. 2016. Endocytic control of cellular signaling at the primary cilium. Trends in Biochemical Sciences 41: 787–797. doi: https://doi.org/10.1016/j.tibs.2016.06.002.

Pusapati, G.V. et al. 2018. CRISPR screens uncover genes that regulate target cell sensitivity to the morphogen Sonic Hedgehog. Developmental Cell 44: 113–129.e118. doi: https://doi.org/10.1016/j.devcel.2017.12.003.

Rack, P.G. et al. 2014. Arhgap36-dependent activation of Gli transcription factors. Proceedings of the National Academy of Sciences of the United States of America 111: 11061–11066. doi: https://doi.org/10.1073/pnas.1322362111, PMID: 25024229.

Radhakrishnan, K., Baltes, J., Creemers, J.W.M. & Schu, P. 2013. Trans-Golgi network morphology and sorting is regulated by prolyl-oligopeptidase-like protein PREPL and the AP-1 complex subunit μ1A. Journal of Cell Science 126: 1155–1163. doi: https://doi.org/10.1242/jcs.116079.

Riobó, N.A., Lu, K., Ai, X., Haines, G.M. & Emerson, C.P. 2006. Phosphoinositide 3-kinase and Akt are essential for Sonic Hedgehog signaling. Proceedings of the National Academy of Sciences of the United States of America 103: 4505–4510. doi: https://doi.org/10.1073/pnas.0504337103, PMID: 16537363.

Rohatgi, R., Milenkovic, L. & Scott, M.P. 2007. Patched1 regulates hedgehog signaling at the primary cilium. Science 317: 372–376. doi: https://doi.org/10.1126/science.1139740, PMID: 17641202.

Scheffzek, K., Ahmadian, M.R. & Wittinghofer, A. 1998. GTPase-activating proteins: helping hands to complement an active site. Trends in Biochemical Sciences 23: 257–262. doi: https://doi.org/10.1016/S0968-0004(98)01224-9.

Schindelin, J. et al. 2012. Fiji: An open-source platform for biological-image analysis. Nature Methods 9: 676–682. doi: https://doi.org/10.1038/nmeth.2019, PMID: 22743772.

Schulz, I. et al. 2002. Modulation of inositol 1,4,5-triphosphate concentration by prolyl endopeptidase inhibition. European Journal of Biochemistry 269: 5813–5820. doi: https://doi.org/10.1046/j.1432-1033.2002.03297.x, PMID: 12444969.

Sepe, M. et al. 2014. Proteolytic control of neurite outgrowth inhibitor NOGO-A by the cAMP/PKA pathway. Proceedings of the National Academy of Sciences of the United States of America 111: 15729–15734. doi: https://doi.org/10.1073/pnas.1410274111.

Stamataki, D., Ulloa, F., Tsoni, S.V., Mynett, A. & Briscoe, J. 2005. A gradient of Gli activity mediates graded Sonic Hedgehog signaling in the neural tube. Genes & Development 19: 626–641. doi: https://doi.org/10.1101/gad.325905.

Stone, D.M. et al. 1996. The tumour-suppressor gene patched encodes a candidate receptor for Sonic hedgehog. Nature 384: 129–134. doi: https://doi.org/10.1038/384129a0.

Stone, D.M. et al. 1999. Characterization of the human Suppressor of fused a negative regulator of the zinc-finger transcription factor Gli. Journal of Cell Science 112: 4437–4448. PMID: 10564661.

Szeltner, Z., Alshafee, I., Juhász, T., Parvari, R. & Polgár, L. 2005. The PREPL A protein, a new member of the prolyl oligopeptidase family, lacking catalytic activity. Cellular and Molecular Life Sciences 62: 2376–2381. doi: https://doi.org/10.1007/s00018-005-5262-5.

Taipale, J. et al. 2000. Effects of oncogenic mutations in Smoothened and Patched can be reversed by cyclopamine. Nature 406: 1005–1009. doi: https://doi.org/10.1038/35023008, PMID: 10984056.

Taipale, J., Cooper, M.K., Maiti, T. & Beachy, P.A. 2002. Patched acts catalytically to suppress the activity of smoothened. Nature 418: 892–897. doi: https://doi.org/10.1038/nature00989, PMID: 12192414.

te Welscher, P. et al. 2002. Progression of vertebrate limb development through SHH-mediated counteraction of GLI3. Science 298: 827–830. doi: https://doi.org/10.1126/science.1075620.

Tempe, D., Casas, M., Karaz, S., Blanchet-Tournier, M.-F. & Concordet, J.-P. 2006. Multisite protein kinase A and glycogen synthase kinase 3 phosphorylation leads to Gli3 ubiquitination by SCF β-TrCP. Molecular and Cellular Biology 26: 4316–4326. doi: https://doi.org/10.1128/mcb.02183-05.

Torres, J.Z., Miller, J.J. & Jackson, P.K. 2009. High-throughput generation of tagged stable cell lines for proteomic analysis. Proteomics 9: 2888–2891. doi: https://doi.org/10.1002/pmic.200800873.

Tukachinsky, H., Lopez, L.V. & Salic, A. 2010. A mechanism for vertebrate Hedgehog signaling: recruitment to cilia and dissociation of SuFu-Gli protein complexes. The Journal of Cell Biology 191: 415–428. doi: https://doi.org/10.1083/jcb.201004108, PMID: 20956384.

Tuson, M. et al. 2011. Protein kinase A acts at the basal body of the primary cilium to prevent Gli2 activation and ventralization of the mouse neural tube. Development 138: 4921–4930. doi: https://doi.org/10.1242/dev.070805, PMID: 22007132.

Vuolo, L., Herrera, A., Torroba, B., Menendez, A. & Pons, S. 2015. Ciliary adenylyl cyclases control the Hedgehog pathway. Journal of Cell Science 128: 2928–2937. doi: https://doi.org/10.1242/jcs.172635.

Wallace, V.A. 1999. Purkinje-cell-derived Sonic hedgehog regulates granule neuron precursor cell proliferation in the developing mouse cerebellum. Current Biology 9: 445–448. doi: https://doi.org/10.1016/S0960-9822(99)80195-X.

Wang, B. & Li, Y. 2006. Evidence for the direct involvement of β-TrCP in Gli3 protein processing. Proceedings of the National Academy of Sciences of the United States of America 103: 33–38. doi: https://doi.org/10.1073/pnas.0509927103.

Wang, C., Pan, Y. & Wang, B. 2010. Suppressor of fused and Spop regulate the stability processing and function of Gli2 and Gli3 full-length activators but not their repressors. Development 137: 2001–2009. doi: https://doi.org/10.1242/dev.052126, PMID: 20463034.

Wang, Y., Zhou, Z., Walsh, C.T. & McMahon, A.P. 2009. Selective translocation of intracellular Smoothened to the primary cilium in response to Hedgehog pathway modulation. Proceedings of the National Academy of Sciences of the United States of America 106: 2623–2628. doi: https://doi.org/10.1073/pnas.0812110106.

Wang, Y.J. et al. 2003. Phosphatidylinositol 4 phosphate regulates targeting of clathrin adaptor AP-1 complexes to the Golgi. Cell 114: 299–310. doi: https://doi.org/10.1016/S0092-8674(03)00603-2, PMID: 12914695.

Wechsler-Reya, R. & Scott, M.P. 2001. The developmental biology of brain tumors. Annual Review of Neuroscience 24: 385–428. doi: https://doi.org/10.1146/annurev.neuro.24.1.385, PMID: 11283316.

Wen, X. et al. 2010. Kinetics of Hedgehog-dependent full-length Gli3 accumulation in primary cilia and subsequent degradation. Molecular and Cellular Biology 30: 1910–1922. doi: https://doi.org/10.1128/mcb.01089-09.

Williams, R.S.B., Eames, M., Ryves, W.J., Viggars, J. & Harwood, A.J. 1999. Loss of a prolyl oligopeptidase confers resistance to lithium by elevation of inositol (1,4,5) trisphosphate. EMBO Journal 18: 2734–2745. doi: https://doi.org/10.1093/emboj/18.10.2734, PMID: 10329620.

Wright, K.J. et al. 2011. An ARL3-UNC119-RP2 GTPase cycle targets myristoylated NPHP3 to the primary cilium. Genes and Development 25: 2347–2360. doi: https://doi.org/10.1101/gad.173443.111, PMID: 22085962.

Zhang, B. et al. 2019. Patched1-ArhGAP36-PKA-Inversin axis determines the ciliary translocation of Smoothened for Sonic Hedgehog pathway activation. Proceedings of the National Academy of Sciences of the United States of America 116: 874–879. doi: https://doi.org/10.1073/pnas.1804042116, PMID: 30598432.

