## Supplementary File 4 for "Multiple domains in ARHGAP36 regulate PKA degradation and Gli activation"

#!/usr/bin/env python
# -*- coding: utf-8 -*-

# ARHGAP36 mutagenesis screen analysis (python 2.7)

# 1. Import libraries of python functions that will be useful for data analysis
import pysam # module for analyzing genomic datasets (manipulation of SAM files etc, samtools/bcftools functionality)
from mock import Mock # allows you to replace parts of your system under test with mock objects and make assertions about how they have been used
import numpy as np # fundamental package for scientific computing with Python.
import csv # implements classes to read and write tabular data in CSV format
import os # allows you to interface with the underlying operating system that Python is running on – be that Windows, Mac or Linux
import pandas as pd # converts CSVs, etc into data frames, useful for data analysis
from os.path import join # concatenates things
from Bio.Seq import Seq # manipulation of DNA/RNA/protein sequences
from collections import OrderedDict # dictionary subclass that remembers order by which elements are inserted

# 2. Define and translate reference sequence (Amplicon produced from wild-type ARHGAP36 Iso2 plasmid – note that only nucleotides 60 – 1608 are CDS)
reference = 'CACGTGAAGGCTGCCGACCCCGGGGGTGGACCATCCTCTAGACTGCCGCTCGAGGCCACCATGAGTGTCTTGGGAGGAGCCCCAGGACACAACCCCGACCGCAGGACGAAGATGGTATCGATACACAGCCTCTCTGAGCTGGAGCGTCTGAAGCTGCAAGAGACTGCTTACCACGAACTCGTGGCCAGACATTTCCTCTCCGAATTCAAACCTGACAGAGCTCTGCCTATTGACCGTCCGAACACCTTGGATAAGTGGTTTCTGATTTTGAGAGGACAGCAGAGGGCTGTATCACACAAGACATTTGGCATTAGCCTGGAAGAGGTCCTGGTGAACGAGTTTACCCGCCGCAAGCATCTTGAACTGACAGCCACGATGCAGGTTGAAGAAGCCACCGGTCAGGCTGCGGGCCGTCGTCGGGGAAACGTGGTGCGAAGGGTGTTTGGCCGCATCCGGCGCTTTTTCAGTCGCAGGCGGAATGAGCCCACCTTGCCCCGGGAGTTCACTCGCCGTGGGCGTCGAGGTGCAGTGTCTGTGGATAGTCTGGCTGAGCTGGAAGACGGAGCCCTGCTGCTGCAGACCCTGCAGCTTTCAAAAATTTCCTTTCCAATTGGCCAACGACTTCTGGGATCCAAAAGGAAGATGAGTCTCAATCCGATTGCGAAACAAATCCCCCAGGTTGTTGAGGCTTGCTGCCAATTCATTGAAAAACATGGCTTAAGCGCAGTGGGGATTTTTACCCTTGAATACTCCGTGCAGCGAGTGCGTCAGCTCCGTGAAGAATTTGATCAAGGTCTGGATGTAGTGCTGGATGACAATCAGAATGTGCATGATGTGGCTGCACTCCTCAAGGAGTTTTTCCGTGACATGAAGGATTCTCTGCTGCCAGATGATCTGTACATGTCATTCCTCCTGACAGCAACTTTAAAGCCCCAGGATCAGCTTTCTGCCCTGCAGTTGCTGGTCTACCTGATGCCACCCTGCCACAGTGATACCCTGGAGCGTCTGCTGAAGGCCCTGCATAAAATCACTGAGAACTGCGAGGACTCAATTGGCATTGATGGACAGTTGGTCCCAGGCAACCGTATGACTTCCACTAACTTGGCCTTGGTGTTTGGATCTGCTCTCCTGAAAAAAGGAAAGTTTGGCAAGAGAGAGTCCAGGAAAACAAAGCTGGGGATTGATCACTATGTTGCTTCTGTCAATGTGGTCCGTGCCATGATTGATAACTGGGATGTCCTCTTCCAGGTGCCTCCCCATATTCAGAGGCAGGTTGCTAAGCGCGTGTGGAAGTCCAGCCCGGAAGCACTTGATTTTATCAGACGCAGGAACTTGAGGAAGATCCAGAGTGCACGCATAAAGATGGAAGAGGATGCACTACTTTCTGATCCAGTGGAAACCTCTGCTGAAGCCCGGGCTGCTGTCCTTGCTCAAAGCAAGCCTTCTGATGAAGGTTCCTCTGAGGAGCCAGCTGTGCCTTCCGGCACTGCCCGTTCCCATGACGATGAGGAAGGAGCGGGTAACCCTCCCATTCCGGAGCAAGACCGCCCATTGCTCCGTGTGCCCCGGGAGAAGGAGGCCAAAACTGGCGTCAGCTACTTCTTTCCTGACCCAGCTTTCTTGTACACCGCGGTTGATCCCGCCACCATGGTGAGCAAGGGCGAGGAGGATAACATGGCCA'

# 3. Define functions to assess reads from sequencing data set:

# 3A. To determine if the read is mapped contiguously to the reference sequence (no indels) (CIGAR string) with paired reads that map to each other and in correct orientation (SAM flags):
def check_read(read):
    if read.cigarstring.count('M') == 1 and read.cigarstring.endswith('M') and read.is_paired and read.flag in (163, 99, 83, 147):
        return True
    else:
        return False

# 3B. To list the read positions (n) in which the read base pair (i) differs from the corresponding base pair (r0) in the reference sequence (ref):
def detect_diff(ref, tex):
    if not ref == tex:
        return [(n, i) for n, (i, r0) in enumerate(zip(tex, ref)) if i != r0]
    else:
        return []

# 3C. To determine the quality score (s) of the variant base pairs in the read:
def detect_diff_score(ref, tex, score):
    if not ref == tex:
        return [(n, i, s) for n, (i, r0, s) in enumerate(zip(tex, ref, score)) if i != r0]
    else:
        return []

# 3D. To determine the nucleotide position of variant base pairs, in the context of the ARHGAP36 CDS:
#	  i. create groupings of all variants' nucleotide positions (s0), all variants' nucleotides, and all variants' quality scores 
#	 ii. to all the nucleotide positions, add the starting nucleotide position of the read in the context of the ARHGAP36 CDS
#	iii. reconstruct the list of (nucleotide position in CDS, nucleotide, quality score) 
def correct_pos(refstart, scores):
    if not scores:
        return []
    s0, s1, s2 = zip(*scores)
    s0 = [i+refstart for i in s0]
    return zip(s0, s1, s2)

# 3E. To generate lists of each read's nucleotide mutations relative to CDS, combine functions 3C and D: 
def detect_var(read):
    scores = detect_diff_score(read.get_reference_sequence(), read.seq, read.query_qualities)
    return correct_pos(read.reference_start, scores)

# 3F. To generate lists of each read's nucleotide mutations relative to CDS, combine functions 3C and D 
#		checking first that the read has mapped to the reference sequence defined in #2.
def detect_var_check_ref(read, reference):
    scores = detect_diff_score(read.get_reference_sequence(), read.seq, read.query_qualities)
    assert read.get_reference_sequence().upper() == reference[read.reference_start:read.reference_start+read.alen]
    return correct_pos(read.reference_start, scores)

# 3G. Identify nucleotides in read which meet quality score threshold.
#		.tolist() necessary to return read quality scores as list of scalars
def ret_positions_above_quality(read, thres):
	return [i for i, ii in zip(read.get_reference_positions(), read.query_qualities.tolist()) if ii > thres]

# 3H. Identify reads which contain interior stop codons:
#	  i. Define _str2array and _array2str as functions to interchange between strings and arrays.
def _str2arr(st):
    assert isinstance(st, str)
    return np.array(list(st))

def _arr2str(ar):
    assert isinstance(ar, np.ndarray)
    return ''.join(ar.tolist())
    
#	 ii. Convert the reference nt sequence into an array (refa)
#	iii. Zip together the list of variant nucleotides in the reads (nuc) with their positions (pos).
#	 iv. Using the list of variant nucleotide positions (p0), replace the wt nucleotide in the reference array at position with the variant nucleotide (n0).
def construct_reads(reference, pos, nuc):
    refa = _str2arr(reference)
    for p0, n0 in zip(pos, nuc):
        refa[p0] = n0
    return _arr2str(refa)

#	  v. Define the CDS section of the reference sequence (bp 60 – 1608) and scan each codon (bp triplet) for stop codons (TAG, TGA, TAA).
def is_stopped(reads):
    tr = reads[60:1608]
    return [tr[i:i+3] for i in range(0, len(tr), 3) if "TAG" in tr[i:i+3] or "TGA" in tr[i:i+3] or "TAA" in tr[i:i+3]]

# 4. Prepare output folder for all outputs of script (one for each sample):

def _make_output(output):
    try:
        os.makedirs(output)
    except:
        pass

# 5. Combine the above functions to determine nucleotide mutations and measure each base pair's mutation frequencies. 

def compiled(input_sam, SCORE_THRES, MAX_MUT, output):
    
    # Collect flags for reads in an ordered dictionary: 
    flag = OrderedDict()
    flag['first_pass_failed'] = 0
    flag['bad_quality'] = 0
    flag['no_mutation'] = 0
    flag['too_many_mutations'] = 0
    flag['stop_codon_middle'] = 0
    flag['bad_quality2'] = 0
    
    # Make output directory, mock object, and empty array the length of the reference sequence (refarr):
    _make_output(output)
    prev = Mock()
    refarr = np.zeros((len(reference)), dtype=np.uint32)

    # Open SAM file as a BAM file ("rb")
    samfile = pysam.AlignmentFile(input_sam, "rb")
    prev.qname = None
    
    # Open a csv file in the output folder for writing ('w'), label it f0, and create a writing object f0w that converts the f0 data into delimited strings (for writing):
    with open(join(output, 'filtered.csv'), 'w') as f0:
        f0w = csv.writer(f0)
        # Number each read of all mapped reads in the opened SAM file (sorted by name). For each read: 
        for num, read in enumerate(samfile.fetch()): 
            # 1. Assert that it passes the quality criteria outlined in the check_read function. If it does (returns a value of 'True'), then continue with the function.
            if not check_read(read):
                flag['first_pass_failed'] += 1
                continue    
            # 2. Ensure that only every other read is processed. Since the SAM file is sorted, this means that read pairs are processed one at a time.
            if read.qname == prev.qname:
                # Ensure that the reads in each pair have bases with good quality.
                positions = ret_positions_above_quality(read, SCORE_THRES) + ret_positions_above_quality(prev, SCORE_THRES)
                if not positions:
                    flag['bad_quality'] += 1
                    continue
                # List the nucleotide positions (pos), identity (nuc) and quality scores (qscore) for each variant detected in the two paired reads.
                scores = detect_var_check_ref(read, reference)
                scores2 = detect_var_check_ref(prev, reference)
                scores.extend(scores2)
                if not scores:
                    flag['no_mutation'] += 1
                if scores:
                    pos, nuc, qscore = zip(*scores)
                # Exclude reads with too many mutations
                if len(pos) >= MAX_MUT:
                    flag['too_many_mutations'] += 1
                    continue
                # Exclude reads with interior stopped codons
                if is_stopped(construct_reads(reference, pos, nuc)):
                    flag['stop_codon_middle'] += 1
                    continue
                # Tabulate nucleotide coverage: in the reference sequence array, add up the number of times a position is covered with good quality in the assessed read pair.
                refarr[np.unique(positions)] += 1
                
                # In the filtered.csv table (f0w), input the nucleotide positions (pos) and identifies (nuc) of the variants detected in the read pair 
                # 1. Create a mask (boo) based on if the corresponding quality score (qscore; converted to integers) exceeds the user-defined score threshold.
                boo = np.array(map(int, qscore)) > SCORE_THRES
                # 2. Apply mask to arrays of the variant nucleotide positions (fp) and variant nucleotide identities (fn) to produce arrays describing variants passing qualty score threshold.
                fp, fn = np.array(pos)[boo].tolist(), np.array(nuc)[boo].tolist()
                if not fp:
                    flag['bad_quality2'] += 1
                if fp:
                    f0w.writerow(fp)
                    f0w.writerow(fn)
            prev = read
    samfile.close()
    np.save(join(output, 'refarr'), refarr)
    print 'number of all reads: {0}'.format(num)
    for key, value in flag.iteritems():
        print key, value
    
    # Calculate nucleotide mutation frequencies.
    # Generate empty array to add number of times a nucleotide position is mutated in the population.
    mutarr = np.zeros((len(reference)), dtype=np.uint32)
    # Open the filtered.csv file written in Step 5 as a file for reading ('r'), label it r0, and create a reading object r1:
    with open(join(output, 'filtered.csv'), 'r') as r0:
        r1 = csv.reader(r0)
        # Restrict analysis to odd rows, which contains the positions of variant nucleotides identified in entire population:
        for num, tex in enumerate(r1):
            if not num % 2 == 0:
                continue
            # Create a set object (mt_positions; unordered, no duplicates) consisting of all the variant nucleotide positions.
            # In the empty mutarray, add up the number of times a position (mt_pos) is variant:
            mt_positions = set(map(int, tex))
            for mt_pos in mt_positions:
                mutarr[mt_pos] += 1
    # Print out the coverage and number of mutated reads for each nucletodie in the CDS (summary.csv)
    df = pd.DataFrame(dict(coverage=refarr[60:1608], mutation=mutarr[60:1608]), index=list(reference)[60:1608])
    df.to_csv(join(output, 'summary.csv'))

# 6. Determine the amino acid mutations resulting from identified nucleotide mutations (pesudo-CDS).

def translate_reads(output):
    aaref = str(Seq(reference).translate())[20:536]
    s0 = 0

    # Open the filtered.csv file written in the section above as a file for reading ('r'), label it r0 and create a reading object r1:
    # Create a aaseq.txt as a writing object ('w'):
    with open(join(output, 'filtered.csv'), 'r') as r0, open(join(output, 'aaseq.txt'), 'w') as w0:
        r1 = csv.reader(r0)
         # In the filtered.csv file, define every odd row as nucleotide position and every even row as residues:
        for num, tex in enumerate(r1):
            if num % 2 == 0:
                positions = tex
            else:
                residues = tex
    
                # Convert the reference nucleotide sequence into a list(seq).
                seq = list(reference[:])
                #  For each read, create a pesudo-CDS which harbors read's nucleotide mutations (res) at the corresponding base pair (pos).
                for pos, res in zip(positions, residues):
                    seq[int(pos)] = res
    
                # Translate these new, mutant pseudo-CDS. If the amino acid sequence is the same, list as s0, if not write in the aaseq.txt file.
                aaseq = str(Seq(''.join(seq)).translate())[20:536]
                if aaseq == aaref:
                    s0 += 1
                else:    
                	w0.write(aaseq+'\n')
    print 'no mutation in terms of aa: {0}'.format(s0)

# 7. Determine each residue's mutation frequencies.

# Define objects describing WT protein sequence and WT protein length
aa_ref = 'MSVLGGAPGHNPDRRTKMVSIHSLSELERLKLQETAYHELVARHFLSEFKPDRALPIDRPNTLDKWFLILRGQQRAVSHKTFGISLEEVLVNEFTRRKHLELTATMQVEEATGQAAGRRRGNVVRRVFGRIRRFFSRRRNEPTLPREFTRRGRRGAVSVDSLAELEDGALLLQTLQLSKISFPIGQRLLGSKRKMSLNPIAKQIPQVVEACCQFIEKHGLSAVGIFTLEYSVQRVRQLREEFDQGLDVVLDDNQNVHDVAALLKEFFRDMKDSLLPDDLYMSFLLTATLKPQDQLSALQLLVYLMPPCHSDTLERLLKALHKITENCEDSIGIDGQLVPGNRMTSTNLALVFGSALLKKGKFGKRESRKTKLGIDHYVASVNVVRAMIDNWDVLFQVPPHIQRQVAKRVWKSSPEALDFIRRRNLRKIQSARIKMEEDALLSDPVETSAEARAAVLAQSKPSDEGSSEEPAVPSGTARSHDDEEGAGNPPIPEQDRPLLRVPREKEAKTGVSYFFP'
aa_len = len(aa_ref)

# Create an array estimating read coverage for each codon (aa_refarr) by averaging the read coverage of each nucleotide in that codon as determined in refarr.
def aa_coverage(output):
	refarr = np.load(join(output, 'refarr.npy'))[60:1608]
	aa_refarr = []
	nums = range(0, len(refarr)+3, 3)
	for n0, n1 in zip(nums[:-1], nums[1:]):
		aa_refarr.append(np.mean(refarr[n0:n1]))
	pd.DataFrame(aa_refarr, index=range(aa_len)).to_csv(join(output,'aa_refarr.csv'))
			
# Define function to identify the residues (num) containing variant residues in each translated pseudo-CDS.
def detect_diff_pos(tex0, tex1):
    return [num for num, (i0, i1) in enumerate(zip(tex0, tex1)) if not i0 == i1]

# Count the number of reads encoding each amino acid mutation for each residue.
def count_aa_mut(output):
    q0 = 0
    t0 = 0

    # Create an empty table (aa_mut) the length of the ARHGAP36 protein sequence with columns for each possible amino acid residue.
    aa_list = ['A','C','D','E','F','G','H','I','K','L','M','N','P','Q','R','S','T','V','W','Y','*']
    aa_mut = pd.DataFrame(np.zeros((21, aa_len), dtype=np.uint32), index=aa_list)

    # Open the aaseq.txt file written in the section above as a file for reading ('r') and label it f:
    with open(join(output, 'aaseq.txt'), 'r') as f:
        # Number each row aka pseudo-CDS (line).
        for n, line in enumerate(f):
            line = line[:-1]
            # Identify the residues which contain mutations (diff_pos).
            diff_pos = detect_diff_pos(aa_ref, line)
            # Identify the amino acid mutation identities (diff_pos_aa) in the variant positions (dp).
            diff_pos_aa = [line[dp] for dp in diff_pos]
            # Count the number of pseudo-CDS that have stop codons.
            if '*' in diff_pos_aa:
                t0 += 1
            else:
            	q0 += 1
            # Tally the variant residue positions (pos) and identities (aa) in aa_mut:
            for (pos, aa) in zip(diff_pos, diff_pos_aa):
                aa_mut[pos][aa] += 1
    aa_mut.to_csv(join(output, 'aa_mut.csv'))
    print 'number of reads with stop mutations: {0}'.format(t0)
    print 'number of reads with non-stopped mutations: {0}'.format(q0)

# Functions for running on command line 
def _parse_command_line_args():
    import argparse
    parser = argparse.ArgumentParser()
    parser.add_argument(
        '--input', help='supply path to sam files',
    )
    parser.add_argument(
        '--score', type=int, help='threshold for scores (e.g. 19, 31, 35)', default=19,
    )
    parser.add_argument(
        '--max_mut', type=int, help='a maximum number of mutations per read pairs (recommend < 11 for calculation cost later)', default=10,
    )
    parser.add_argument(
            '--output', help='directory name for output', default='output')
    return parser.parse_args()

def _main():
    args = _parse_command_line_args()
    print args.input
    print args.score
    print args.max_mut
    print args.output
    compiled(args.input, int(args.score), int(args.max_mut), args.output)
    print "compiled done"
    translate_reads(args.output)
    print "translate_reads done"
    aa_coverage(args.output)
    print "aa_coverage done"
    count_aa_mut(args.output)
    print "count_aa_mut done"

if __name__ == "__main__":
    _main()
    print "script complete"
